# Recycling of trans-Golgi SNAREs is essential for apoplastic effector secretion and effective pathogenicity of *Magnaporthe oryzae*

**DOI:** 10.1101/2024.08.02.606313

**Authors:** Lili Lin, Qiuqiu Wu, Shuang Wang, Qing Gong, Xiuwei Huang, Yakubu Saddeeq Abubakar, Yue Liu, Jiaying Cao, Jiexiong Hu, Zonghua Wang, Guodong Lu, Wenhui Zheng

## Abstract

Vesicle transport is an essential process that mediates the growth, development and virulence of pathogenic fungi. However, the intricate mechanisms underlying how vesicle transport regulates the secretion of effector proteins remain to be fully elucidated. Here, we unveiled a novel pathway in which retromer and trans-Golgi (TGN) SNARE proteins co-regulate the proper secretion of apoplastic effectors in the rice blast fungus *Magnaporthe oryzae.* A TGN-associated SNARE complex consisting of MoSnc1, MoTlg1, MoTlg2, and MoVti1 was found to be essential for growth, development and pathogenicity in the fungus. Moreover, the TGN-associated SNARE complex is indispensable for accurate secretion of apoplastic effectors. Furthermore, we have elucidated that the dynamin-like protein MoVps1, an upstream regulator of the retromer complex, regulates the fission of MoVps35-coated vesicle and the proper localization of the TGN-associated SNARE complex. Additionally, we employed prochlorperazine, which identified as a potent dynamin inhibitor, elicits a developmental response in *M. oryzae* akin to *MoVPS1* disruption, highlighting the pivotal regulatory role of dynamin and its potential as a therapeutic target for rice blast disease management. In conclusion, the study uncovered a specific mechanism by which MoVps1 and the retromer complex regulate the positioning of TGN-associated SNARE proteins to effectively promote effector secretion. It provides a deeper understanding of the molecular mechanisms of effector secretion in fungi and underscores the importance of vesicle transport in fungal pathogenesis.

**Importance:** Vesicle transport is essential for pathogenic fungi as it controls the secretion of effectors that modulate interactions with the host and infection processes. The detailed mechanisms of effector secretion via vesicular pathways in these fungi are not yet fully understood. In this study, we have discovered a new regulatory pathway involving the retromer complex and trans-Golgi SNARE proteins that is critical for the proper secretion of apoplast effectors in *M. oryzae.* We have identified an important TGN-associated SNARE complex, consisting of MoSnc1, MoTlg1, MoTlg2 and MoVti1, which is required for the development and pathogenicity of *M. oryzae.* Our results emphasize the importance of this SNARE complex for the precise secretion of effectors into the apoplast, a key step in pathogenesis. Additionally, we demonstrated that the dynamin-like protein MoVps1, a protein acting upstream of the retromer complex, is vital for the correct localization of the TGN-associated SNARE complex. Furthermore, our research underscores the critical regulatory role of dynamin in *M. oryzae* pathogenesis, with prochlorperazine serving as an inhibitor that mimics the phenotypic effects of MoVps1 disruption, thereby highlighting its potential as a biopesticide candidate for rice blast disease management. Our study has uncovered a specific regulatory mechanism in which MoVps1 and the retromer complex control the positioning and function of TGN-associated SNARE proteins, thereby facilitating effector secretion. This work not only advances our understanding of the molecular basis of effector secretion in fungi, but also has implications for the development of novel strategies to control fungal diseases.

## Introduction

Endosomal transport is a cornerstone of cellular function and orchestrates the intricate ballet of molecular and organelle movement that maintains the integrity and functionality of the cell[1, 2]. Central to this system are the membrane-bound endosomes, which play a crucial role in regulating nutrient uptake and protein homeostasis. The process is initiated by endocytosis, whereby internalized molecules are directed to early endosomes, which mature late endosomes, where their cargoes are sorted for subsequent degradation and/or recycling [3, 4]. The trans-Golgi network (TGN) is a master regulator that directs proteins to their final destinations, ensuring the smooth operation of the cellular machinery [5, 6]. The selective packaging and sorting signals of the TGN are essential for precise protein delivery, which is essential for efficient cell function[7, 8]. In Addition, endosomal transport regulates signal transduction and controls the fate of signaling molecules in a responsive cellular environment[9]. However, the complicated nature of endosomal transport also poses a complex challenge in the context of pathogen pathogenicity.

*Magnaporthe oryzae* is a fungal pathogen responsible for the devastating rice blast disease [10]. The specificity and efficiency of effector secretion are key to the virulence for pathogens to undermine host immunity [11–14]. The effectors of *M. oryzae* can be divided into two categories according to the different secretion pathways [15]. Firstly, there are cytoplasmic effectors such as Avr-Pita and Pwl2 that accumulate in the biotrophic interfacial complex (BIC), a structure that includes exocyst components and a novel t-SNARE (Soluble N-ethylmaleimide-sensitive Factor Attachment protein Receptor) secretory pathway [15]. Another pathway involves apoplastic effectors such as Bas4 and Slp1, which are secreted via the conventional ER-Golgi-dependent pathway and accumulate between the fungal cell wall and the rice plasma membrane [16]. Therefore, the functional diversity of effectors largely depends on their transport pathways and secretion mechanisms. However, the precise regulatory mechanisms governing the vesicular transport of effectors, particularly through the trans-Golgi network and the retromer-regulated SNARE pathways, are not fully understood.

Effectors are secreted via a conserved pathway involving the TGN and SNARE proteins which are essential for the fusion of vesicles with their target membranes [17]. SNARE proteins are important for vesicle trafficking and facilitate the fusion of the vesicles with their target membranes, an important process for the transport of cellular cargoes [18]. They are a conserved superfamily characterized by SNARE motifs and a single transmembrane domain or lipid modification motif [19–21]. The precise assembly of these proteins into a four-helix bundle is essential for driving membrane fusion, a process that mediates the proper targeting and release of cellular cargoes [22, 23]. Recycling of SNARE proteins is facilitated by the N-Ethylmaleimide Sensitive Factor (NSF), an ATPase enzyme that disassembles the SNARE complex, thereby preparing the components for further rounds of the fusion events [17]. This highly regulated and dynamic process of vesicle transport and membrane fusion is essential for the proper functioning and survival of fungi.The understanding of the specificity of SNARE-mediated fusion events has been informed by genetic studies that have identified the localizations of the various SNARE components [19, 20]. For instance, the yeast *Saccharomy cescerevisiae* possesses a unique set of SNAREs, including the syntaxins which are integral to the vesicular transport process [19]. The specificity of these proteins is thought to be encoded in the pattern of their interactions, with each organelle potentially recognized by a unique SNARE or a specific set of SNAREs. Recent studies have also highlighted the role of additional factors, such as Rab/Ypt GTPases and peripheral membrane proteins, in the initial docking events that precede SNARE-mediated membrane fusion[24, 25]. These findings suggest a complex interplay between various proteins in determining the selectivity and efficiency of vesicle fusion.

MoSnc1 is crucial for cargo sorting and effector secretion in *M. oryzae*. Our prior research has delineated the role of MoRab7/Retromer/MoSnc1 sorting machinery in effector secretion during the biotrophic and invasive phases of the rice blast fungus [13]. To further elucidate the intricate regulatory interplay between the retromer complex and SNARE components, we employed liquid chromatography–tandem mass spectrometry (LC–MS/MS) to ascertain the binding partners of MoSnc1. The results showed that the MoTlg2 (Qa-SNARE), MoVti1 (Qb-SNARE), and MoTlg1 (Qc-SNARE) were found to be highly abundant and all associated with TGN. We further elucidated their functional significance in the regulation of vegetative growth, conidiation, and virulence in *M. oryzae.* In addition, we provided evidence demonstrating the indispensable role of the TGN-associated SNARE proteins in the delivery of apoplastic effectors and that this role relies on the synergistic functions of the dynamin-like GTPase MoVps1 and the retromer core component MoVps35. Moreover, we identified prochlorperazine, which is an inhibitor of dynamin, as a potent inhibitor that significantly impacts *M. oryzae* virulence. Taken together, our study established a new regulation pathway involving MoVps1/retromer/SNARE and provided novel insights into how apoplastic effectors are secreted and regulated during infection.

## Results

### MoSnc1 forms a complex with MoTlg2, MoVti1, and MoTlg1 in *M. oryzae*

The protein MoSnc1 was previously identified as a key player in *M. oryzae* pathogenicity and effector secretion [13, 15], but its specific regulation mechanism requires further studies. SNARE proteins always function as complexes with other Qa-, Qb-, Qc-, or R-SNARE proteins to facilitate membrane fusion [26]. To further unveil the functional mechanisms of MoSnc1 in *M. oryzae*, we immunoprecipitated GFP-MoSnc1 and performed liquid chromatography-tandem mass spectrometry (LC-MS/MS) to identify the SNARE proteins that potentially associate with MoSnc1. A total 9 SNARE proteins were identified, among which MoTlg2 (Qa-SNARE), MoVti1 (Qb-SNARE), and MoTlg1 (Qc-SNARE) were found (Fig. 1A), orthologs of these proteins in yeast are known to be primarily localized at the TGN, where they coalesce into functional complexes[8, 27]. Despite this knowledge, the potential assembly of analogous complexes involving these SNARE proteins in *M. oryzae* has yet to be determined. To ascertain whether these form a functional complex in *M. oryzae*, we firstly used AlphaFold2 to predicate the interactions among these proteins. The results showed that the MoSnc1(R-SNARE), MoTlg2 (Qa-SNARE), MoVti1 (Qb-SNARE) and MoTlg1 (Qc-SNARE) form a typical four-helix SNARE structure of the complex, which displayed an “pTM and ipTM” value of 0.563 (Fig. 1B). Subsequent to our structural predictions, yeast two-hybrid (Y2H) assays were conducted to validate the interactions among the identified SNARE proteins (Fig. S1). Interestingly, The assays revealed that MoSnc1, an R-SNARE, along with MoVti1 and MoTlg1, both Qb- and Qc-SNAREs respectively, exhibit direct physical interactions with the Qa-SNARE protein MoTlg2. Notably, no interactions were detected among the other SNARE proteins, suggesting a potential scaffolding or recruiting role for the MoTlg2 in the assembly of the SNARE complex.

**Fig 1.**
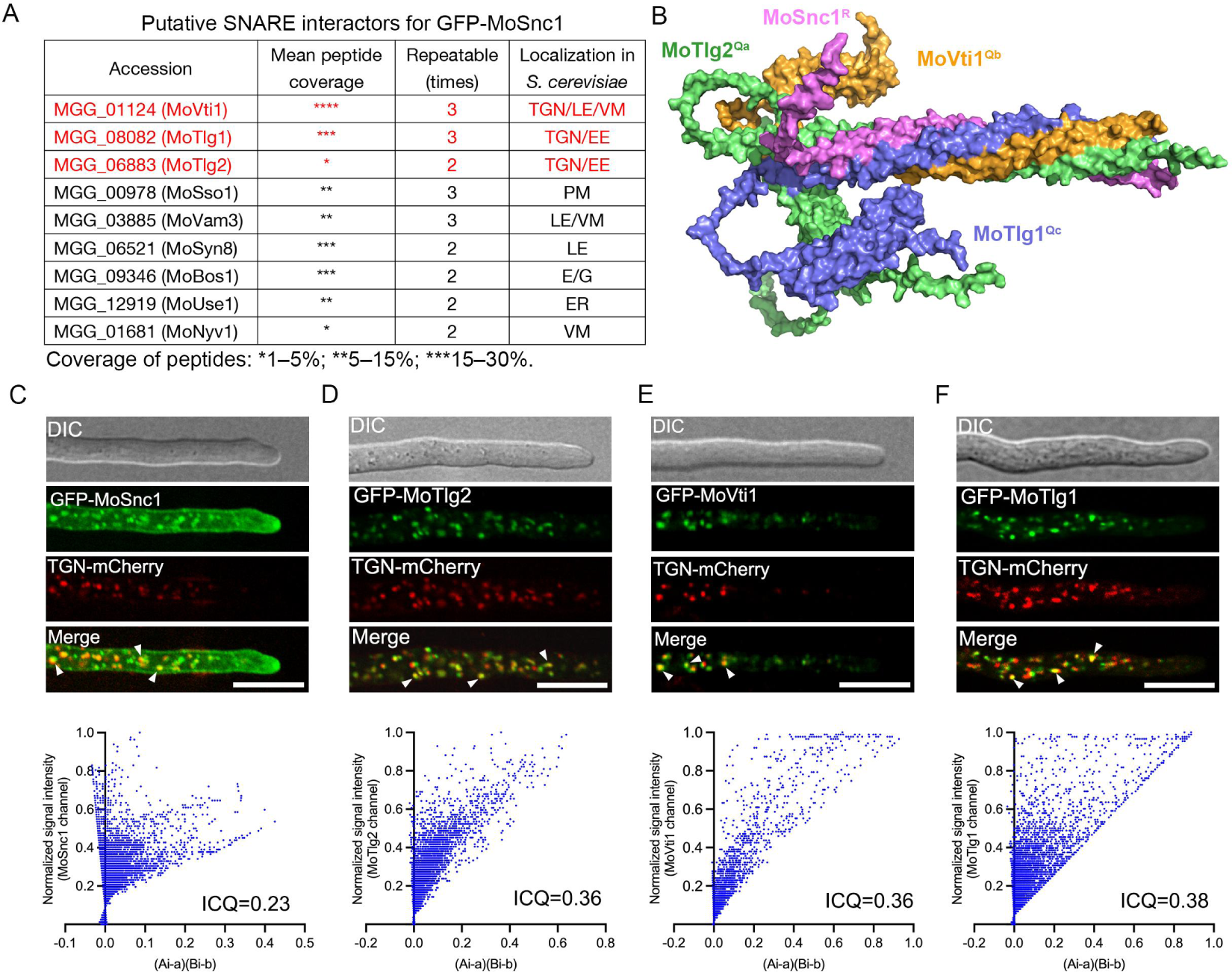
A set of TGN-associated SNARE protein complexes has been identified in the rice blast fungus. A. Liquid chromatography-tandem mass spectrometry (LC-MS/MS) unveiled nine putative GFP-MoSnc1-interacting SNARE proteins in *M. oryzae*. Single, double, and triple asterisks represent the degree of peptide coverage. “TGN” represents tran-Golgi network, “LE” represents late endosome, “EE” represents early endosome, “VM” represents vacuole membrane, “PM” represents plasma membranes, “ER” represents endoplasmic reticulum. B. An AlphaFold-generated model depicting the complex formed by MoSnc1, MoTlg1, MoVti1, and MoTlg2, which displayed an “pTM and ipTM” value of 0.563. C-F. The strains expressingGFP-MoSnc1(C), GFP-MoTlg2 (D), GFP-MoVti1 (E), and GFP-MoTlg1 (F) were observed to exhibit partial co-localization with the TGN marker (MoKex2-mCherry) in the vegetative hyphae of *M. oryzae*. Scale bars=10 µm. Arrows indicate the colocalization of Green and mCherry fluorescence signal. The accompanying graph illustrates the application of Li’s intensity correlation coefficient (ICQ), a metric employed for gauging the extent of colocalization. The terms ‘Ai’ and ‘a’ represent the current and average fluorescence intensities of the SNARE proteins channel, respectively, while ‘Bi’ and ‘b’ correspond to the respective values for the TGN-mCherry channel. A high degree of colocalization is indicated by a pixel cloud distribution towards the right side of the plot. The ICQ scale spans from −0.5, denoting exclusion, to 0.5, signifying complete colocalization.

In addition, the dynamic distribution of these TGN SNAREs was characterized in the vegetative hyphae and invasive hyphae of *M. oryzae* was characterized. We found that MoSnc1 and MoVti1 localized to the plasma and vacuole membranes, respectively; however, all the four SNAREs, including MoTlg1 and MoTlg2, consistently show a distinct punctate localization pattern (Fig. S2). To investigate the association of these SNARE proteins with the TGN, we used the TGN marker MoKex2-mCherry to check for possible colocalization with the SNAREs. Live-cell imaging demonstrated that all the four SNARE proteins exhibited partial co-localization with the MoKex2-mCherry (Fig. 1C-F). Using Li’s correlation analysis, we found that both the cloud shape and the data for MoSnc1 (ICQ=0.23), MoTlg2 (ICQ=0.36), MoVti1 (ICQ=0.36), and MoTlg1 (ICQ=0.38) are consistent with the partial colocalization of the TGN marker. These observations suggest that MoSnc1, MoTlg1, MoTlg2, and MoVti1 could be involved in vesicle transport and fusion by forming a complex at the TGN, thereby contributing to the TGN-mediated vesicle trafficking of *M. oryzae*.

### MoTlg1 and MoTlg2 are important for vegetative growth and conidiation of *M. oryzae*

To further analyze the roles of MoTlg1 and MoTlg2 in *M. oryzae*, we initially attempted to generate knockout mutants in the 70-15 strain background by replacing the open reading frames (ORFs) of *MoTLG1* and *MoTLG2* with a hygromycin resistance gene (HPH). Despite multiple attempts, we were unable to obtain a knockout mutant for *MoTLG2* in the 70-15 background. Consequently, we switched to the Guy11 strain background, where we successfully generated the Δ*Motlg2* knockout mutant. The resulting mutants Δ*Motlg1* and Δ*Motlg2* were confirmed by Southern blot analysis (Fig. S3). To gain insights into the contributions of MoTlg1 and MoTlg2 in the vegetative development of *M. oryzae*, the Δ*Motlg1* and Δ*Motlg2* mutant strains were cultured on different types of nutrient-sufficient and nutrient-deficient culture media including CM, PA, CMII and RBM. Phenotypic characterizations of the mutants on these media revealed that both Δ*Motlg1* and Δ*Motlg2* mutants exhibited a significant decrease in vegetative growth compared to the wild-type and complemented strains (Fig. 2A-E). Moreover, to unravel the potential roles of MoTlg1 and MoTlg2 in asexual reproduction in the rice blast fungus, we monitored conidiophore formation and conidia production. We observed that both Δ*Motlg1* and Δ*Motlg2* mutants produced fewer conidia and conidiophores than the wild-type and complemented strains (Fig. 2E-F), suggesting a defect in asexual reproduction. These findings demonstrate that MoTlg1 and MoTlg2 play essential roles in the vegetative growth and asexual reproduction of *M. oryzae*.

**Fig 2.**
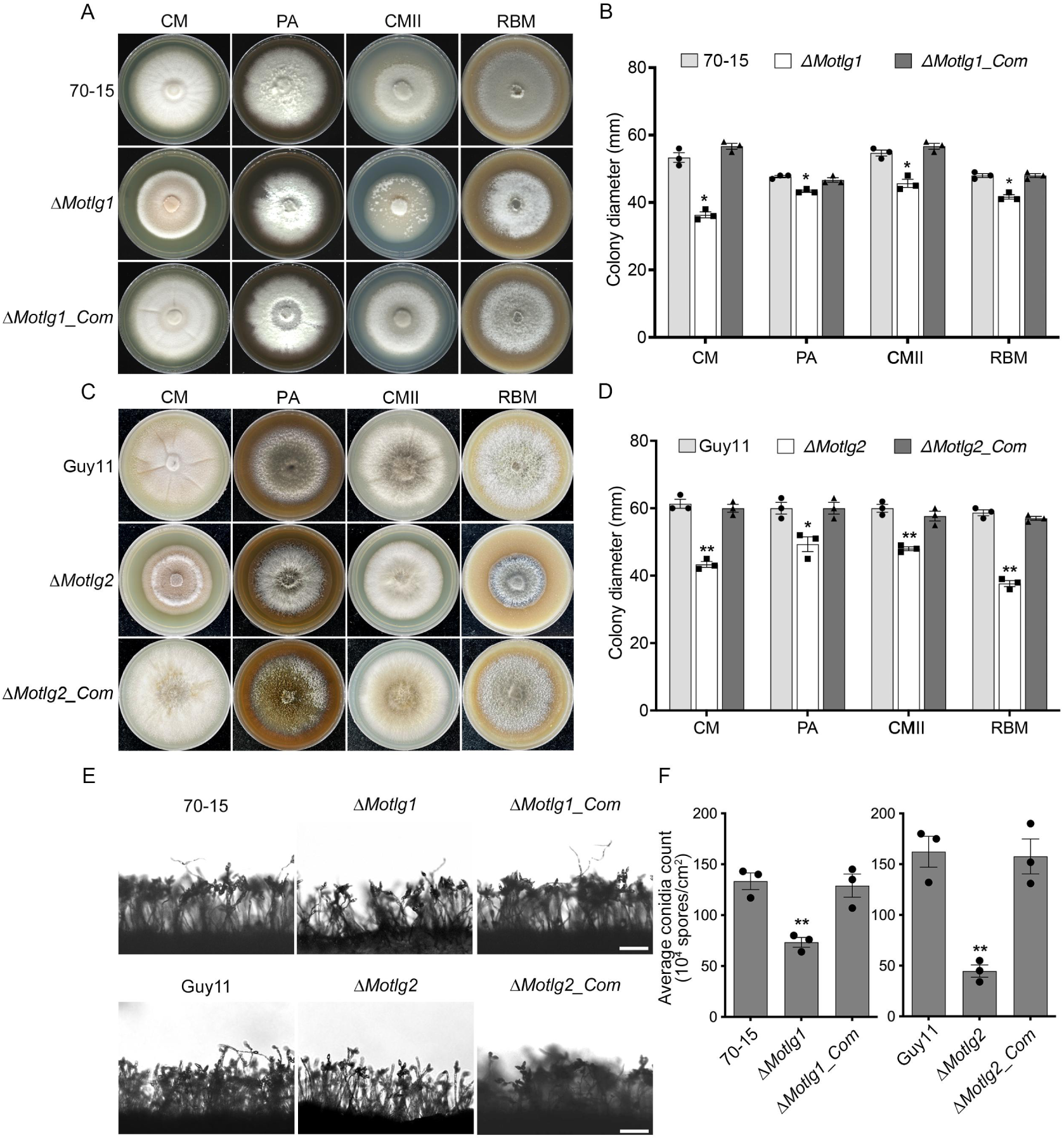
MoTlg1 and MoTlg2 contribute positively to vegetative growth and asexual reproduction of *M. oryzae*. A. Vegetative growths of the WT (wild-type, 70-15), Δ*Motlg1* and Δ*Motlg1_Com* strains cultured on CM, PA, CMII, and RBM for 10 days. B. Statistical analysis of the colony diameters of the indicated strains cultured on CM, PA, CMII and RBM for 10 days. C. Vegetative growths of the WT (wild-type, Guy11), Δ*Motlg2*, and Δ*Motlg2_Com* strains cultured on CM, PA, CMII, and RBM for 10 days. D. Statistical analysis of the colony diameters of the indicated strains cultured on CM, PA, CMII, and RBM for 10 days. E. Conidiophore production potential of the various strains. After vegetative growth phase, cultures of the indicated strains were exposed to 12 h/12 h of light/dark photoperiod for 2 days. Scale bars = 20 µm. F. Analyses of the number of conidia produced by the indicated strains. The error bars represent standard deviation (SD) from three replicates, and single and double asterisks denote significant differences at p< 0.05 and p< 0.01, respectively, based on t-test analysis.

### MoTlg1 and MoTlg2 contribute to the full virulence of *M. oryzae*

To assess the virulence efficiency of the asexual spores produced by Δ*Motlg1* and Δ*Motlg2* mutants, we inoculated both intact and injured barley leaves with spore suspensions (5×10^4^ spores/mL) from the Δ*Motlg1* and Δ*Motlg2* mutants, along with the complemented and wild-type strains. At the same time, leaves of the susceptible rice seedlings (CO39) were spray-inoculated with the spores from the individual strains. Records obtained from these infection assays showed that deletion of *MoTLG1* caused significant reduction in the fungal pathogenicity in both barley and rice inoculated assays, with fewer and smaller lesions observed on the rice leaves compared to the wild-type and complemented strains. The Δ*Motlg2* mutants, however, showed only little reduction in pathogenicity compared to the wild-type and complemented strains (Fig. 3A-B). To explore the potential reasons behind the reduced virulence of Δ*Motlg1* and Δ*Motlg2* mutants, we inoculated rice sheaths with spore suspensions from the various strains, respectively, and monitored their invasive hyphal growths at 36 hours post-inoculation (hpi). Our assessment revealed that Δ*Motlg1* had only 11% incidence of type IV invasive hyphal (IH) growth, which is considerably lower than that of the wild-type (41.3%) and complemented strains (37.3%), respectively. Similarly, Δ*Motlg2* exhibited 31.87% type IV IH growth, in contrast to 67% and 50.3% exhibited by the wild-type and complemented strains, respectively. These results indicated that deletion of *MoTLG1* and *MoTLG2* genes perturbs the ability of the invasive hyphae to proliferate within the plant cells (Fig. 3C-D). We therefore conclude that MoTlg1 and MoTlg2 positively impact *M. oryzae* pathogenicity by promoting the fungal invasive growth in the host.

**Fig 3.**
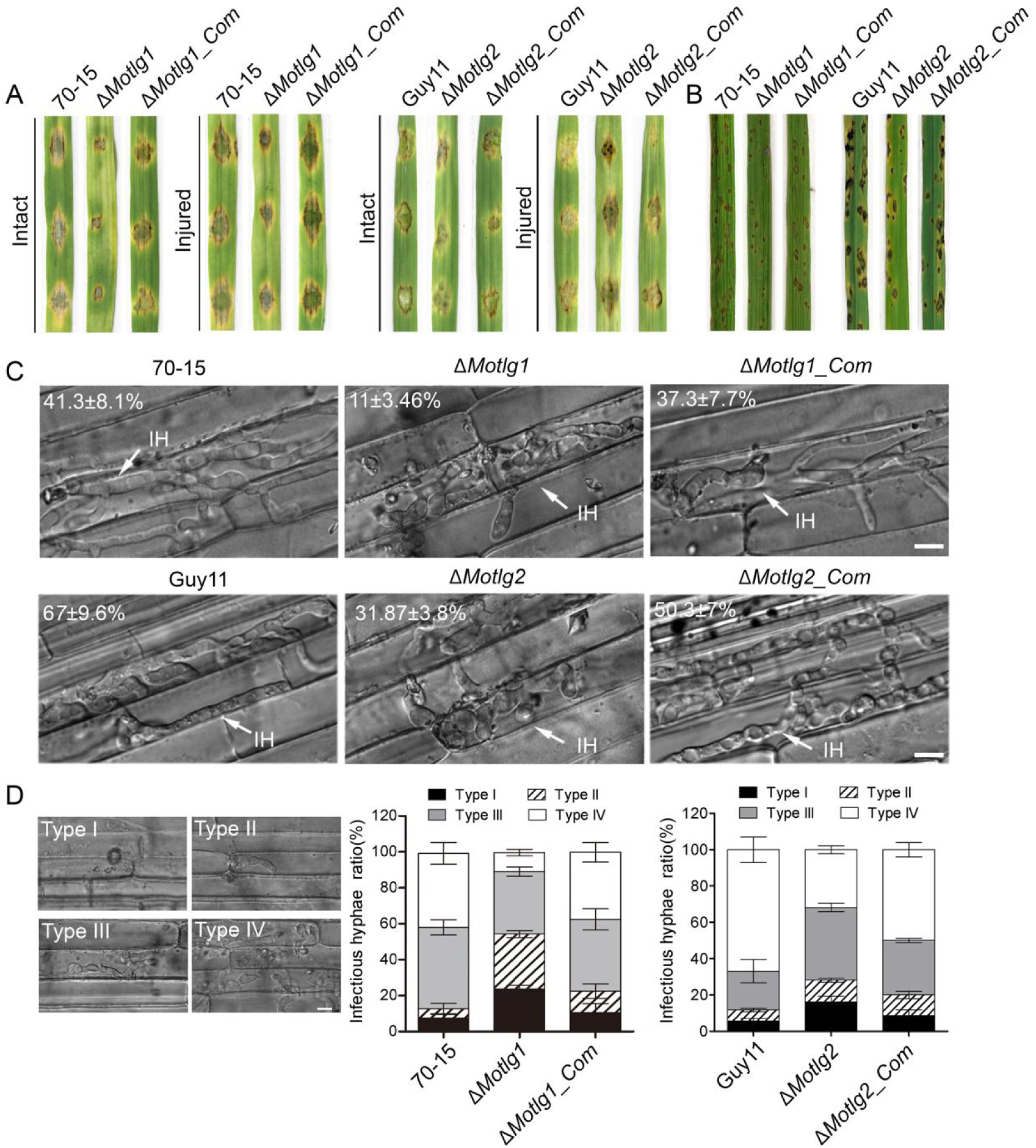
Roles of MoTlg1 and MoTlg2 in the pathogenicity of *M. oryzae*. A. Analysis of the pathogenicity of Δ*Motlg1* and Δ*Motlg2* mutants on intact and injured barley leaves. Conidia suspensions of similar concentrations from the various strains were dropped on the leaves and incubated for 7 days. B. Analysis of the pathogenicity of Δ*Motlg1* and Δ*Motlg2* strains on rice seedlings by spray assay. Conidia suspensions of similar concentrations were sprayed on 3-week-old susceptible CO39 rice seedlings. Disease lesions were observed and photographed at the 7^th^ day post-inoculation (dpi). C. Rice sheath penetration ability of the various strains. The penetration and colonization efficiencies of the wild-type, mutants and complemented strains were analyzed 48 hours post-inoculation of the rice sheaths with spore suspensions. “IH” invasive hyphae, respectively. Scale bars = 10 µm. D. Analyses of the invasive hyphal development of the individual strains. Extent of colonization was classified into four types: type I (appressorium formation), type II (primary invasive hyphae), type III (secondary invasive hyphae), and type IV (extensive hyphal growth). Scale bars = 10 µm. The experiment was repeated three times with three technical replicates each time. For each replicate, a total of 100 infection sites (thus n = 300) were scanned under a microscope.

### *MoVTI1 is* an essential gene that plays vital roles in the growth and pathogenicity of *M. oryzae*

To elucidate the role of *MoVTI1* gene in the growth and pathogenicity of *M. oryzae,* we initially sought to create a knockout mutant of *MoVTI1* using homologous recombination; however, this was not successful after several attempts, prompting us to adopt the use of a Tet-off system for regulated gene expression. Utilizing this system, we effectively generated *MoVTI1-Tet* mutant strains (Fig. S4A). Using confocal microscopy, we examined the impact of doxycycline (DOX) on *MoVTI1* expression in the mutants. We observed a gradual decrease in the intensity of GFP fluorescence with increase in DOX concentrations (Fig. S4B), suggesting the effectiveness of the system in regulating *MoVTI1* expression in a dose-dependent manner in the mutant strains.

The growth of the *MoVTI1-Tet* mutants on CM plates with or without DOX was assessed to unveil the impact of *MoVTI1* silencing on the fungal growth. The mutants exhibited a marked reduction in vegetative growth in the presence of DOX, compared to the untreated controls, supporting the possible lethality of complete deletion of the *MoVTI1* gene in *M. oryzae* (Fig. S4C-D). To assess the impact of *MoVTI1* silencing on the pathogenicity of *M. oryzae,* conidia suspensions from the *MoVTI1-Tet* mutants and the wild-type strain were inoculated on barley leaves, with and without DOX treatment. A significant reduction in disease symptoms was observed in the mutants treated with DOX, compared to the untreated mutants and the wild-type strain (Fig. S4E). Collectively, these results indicate that MoVti1is essential for the survival and plays vital roles in the vegetative growth and pathogenicity of *M. oryzae*.

### TGN-associated SNARE proteins influence apoplast effector section in *M. oryzae*

Our previous findings indicated the significant regulatory role of MoSnc1 in both cytoplasmic and apoplastic effector secretions [13]. We therefore decided to further elucidate the involvement of other TGN-associated SNARE proteins in the effector secretion process of the rice blast fungus. We introduced two fusion vectors harbouring Bas4-GFP and Pwl2-mCherry effector genes, respectively, into the wild-type, Δ*Motlg1*, and Δ*Motlg2* strains to evaluate their subcellular localizations. We observed that disruption of *MoTLG1* and *MoTLG2* to lead Bas4-GFP predominant accumulation of Bas4-GFP in the vacuoles or cytoplasm of the IH, contrary to Pwl2-mCherry localization which had no obvious difference in its localization in the wild-type and the two mutant strains (Fig. 4A and C). This aberrant localization of Bas4 in the mutants suggests a disruption in its secretion pathway, leading to misdirection of the effector protein to the vacuoles, instead of being secreted into the host apoplast. Similarly, treatment of *MoVTI1-Tet* stain with doxycycline let Bas4-mCherry accumulated within the vacuoles while Pwl2-mCherry displayed multiple fluorescent foci, indicating that inactivating *MoVTI1* function results in improper secretion and mis-localization of both cytoplasmic and apoplastic effectors (Fig. 4B-C).

**Fig 4.**
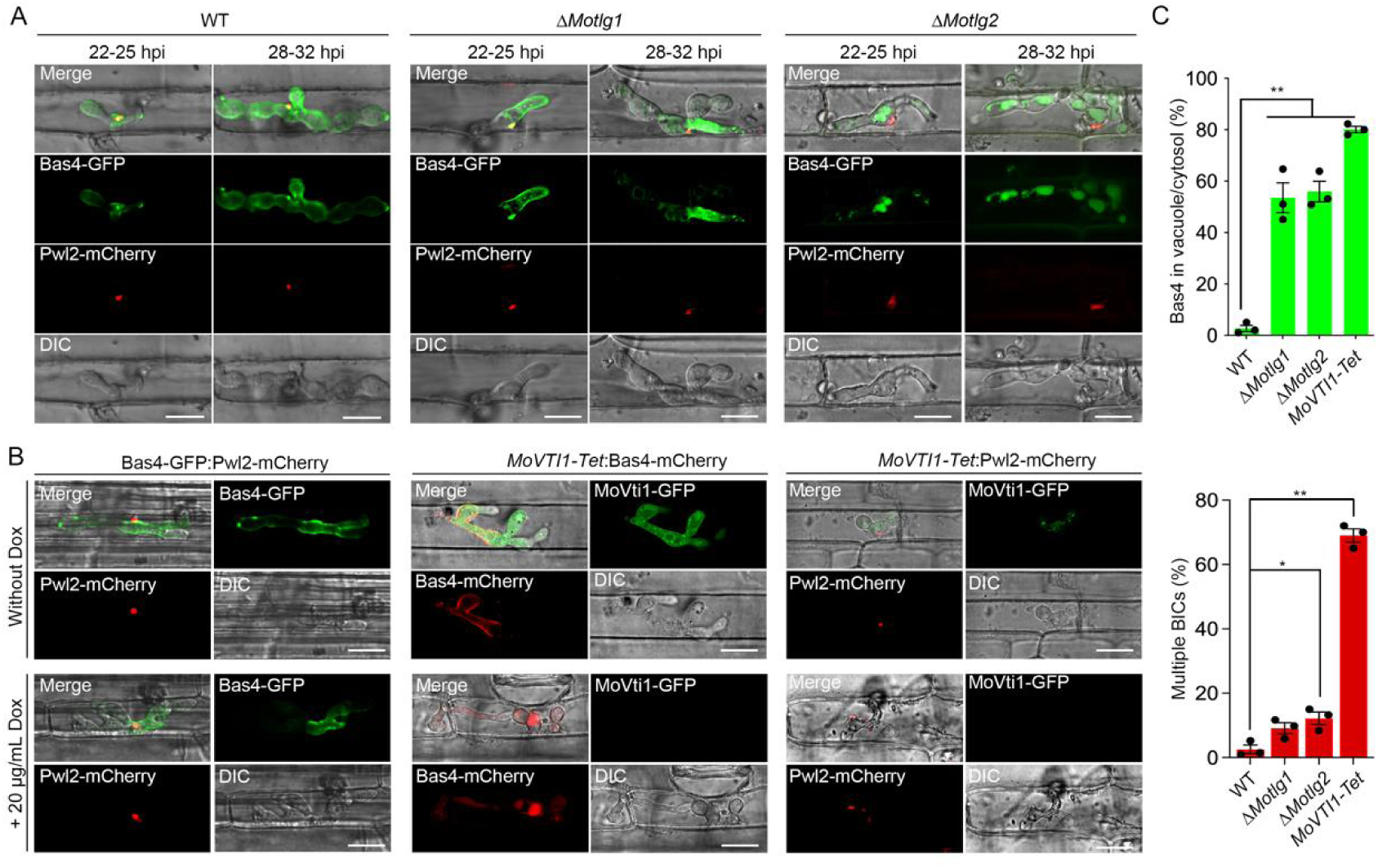
TGN-associated SNARE proteins modulate the classical effector secretory pathway. A. Localization of Bas4-GFP (apoplastic effector) and Pwl2-mCherry (cytoplasmic effector) in the wild-type, Δ*Motlg1* and Δ*Motlg2* strains. The fluorescently tagged Bas4 proteins in the mutant strains were observed to accumulate within the vacuoles of the IH, rather than delineating the periphery of the invasive structures as shown in the WT. B. Relative localizations of two effector proteins in the presence and absence of Dox treatment. Bas4-GFP:Pwl2-mCherry strains as a negative control to verify the Dox won’t affect the localization of this two effector proteins. The two effector proteins were improperly secreted and mis-localized following treatment with 20 μg/mL Dox. The fluorescently tagged Bas4 protein was observed to accumulate within the vacuoles of the IH, rather than delineating the periphery of the invasive structures, whereas the fluorescence of Pwl2 manifests as multiple fluorescent foci within the IH of the mutant, in contrast to its single and concentrated fluorescence at the specialized BIC region in the WT. Scale bars = 10 µm. C. Bar charts illustrating the percentage of Bas4 in vacuole and the IH containing multiple BICs in the WT, Δ*Motlg1*, Δ*Motlg2,* and *MoVTI-Tet* mutant. Error bars represent standard deviation from three replicates. Single and double asterisks denote significant differences at p< 0.05 and p< 0.01, respectively, based on t-test analysis.

Based on the above results, we hypothesized that the TGN-associated SNAREs are involved in the ER-to-Golgi transport pathway, an important pathway for protein sorting and trafficking. Since Brefeldin A (BFA) is known to disrupt protein transport from ER to the Golgi apparatus[28], we analyzed the localization patterns of GFP-MoTlg1, GFP-MoTlg2, GFP-MoVti1, and GFP-MoSnc1 under BFA treatment. We observed that the BFA-treated samples showed marked decrease in the number of punctate structures which accumulated in some compartments that were larger than those in the DMSO-treated controls, whereas BFA-treatment did not affect the plasma membrane and Spitzenkörper localization pattern of GFP-MoSnc1(Fig. S5A-B). These findings suggest that the SNARE complex localizes to Golgi structures during the transport process and may be involved in the specific targeting and secretion of extracellular effector proteins regulated by the trans-Golgi network.

### MoVps35 interacts with and regulates the spatial distribution of TGN-associated SNARE Proteins in *M. oryzae*

Retromer is a multimeric protein complex crucial for regulating the retrieval and recycling of various cargoes, redirecting them away from the degradative pathway and facilitating their delivery to the TGN [29]. Our previous study showed that the core component of the retromer complex MoVps35 recognizes MoSnc1 and facilitates its transport to the plasma membrane [13]. To further explore the intricate molecular interactions and regulatory mechanisms between the TGN-associated SNARE proteins and MoVps35 in *M. oryzae*, we generated strains co-expressing MoVps35-flag and each of GFP-MoTlg1, GFP-MoTlg2 and GFP-MoVti1, respectively, and assayed for possible interaction between each of the protein pairs. Coimmunoprecipitation (Co-IP) assays revealed positive interactions between MoVps35 and each of the SNARE proteins (Fig. 5A). To further validate these results, we performed co-localization analyses between MoVps35 and the respective SNARE proteins. Consistently, GFP-MoTlg1, GFP-MoTlg2 and GFP-MoVti1 showed partial co-localization with MoVps35-mScarlet, as indicated by the white arrows in the zoomed regions in Fig 5B. Using Li’s correlation analysis, both the cloud shape and the data for MoTlg1(ICQ=0.42), MoTlg2 (ICQ=0.34), and MoVti1 (ICQ=0.42) are consistent with the partial colocalization of the proteins with MoVps35(Fig. 5C). To assess the role of MoVps35 in regulating the correct localizations of the TGN-associated SNARE proteins, we performed the fluorescence confocal microscopy imaging assays, and found that loss of *MoVPS35* cause all of the GFP-tagged SNARE proteins to be misdirected to the vacuolar membrane but not the punctate structures (Fig. 5D). We therefore infer that the retromer complex orchestrates the precise localization of the SNARE proteins that are associated with the TGN, ensuring their appropriate placement within this crucial subcellular localization.

**Fig 5.**
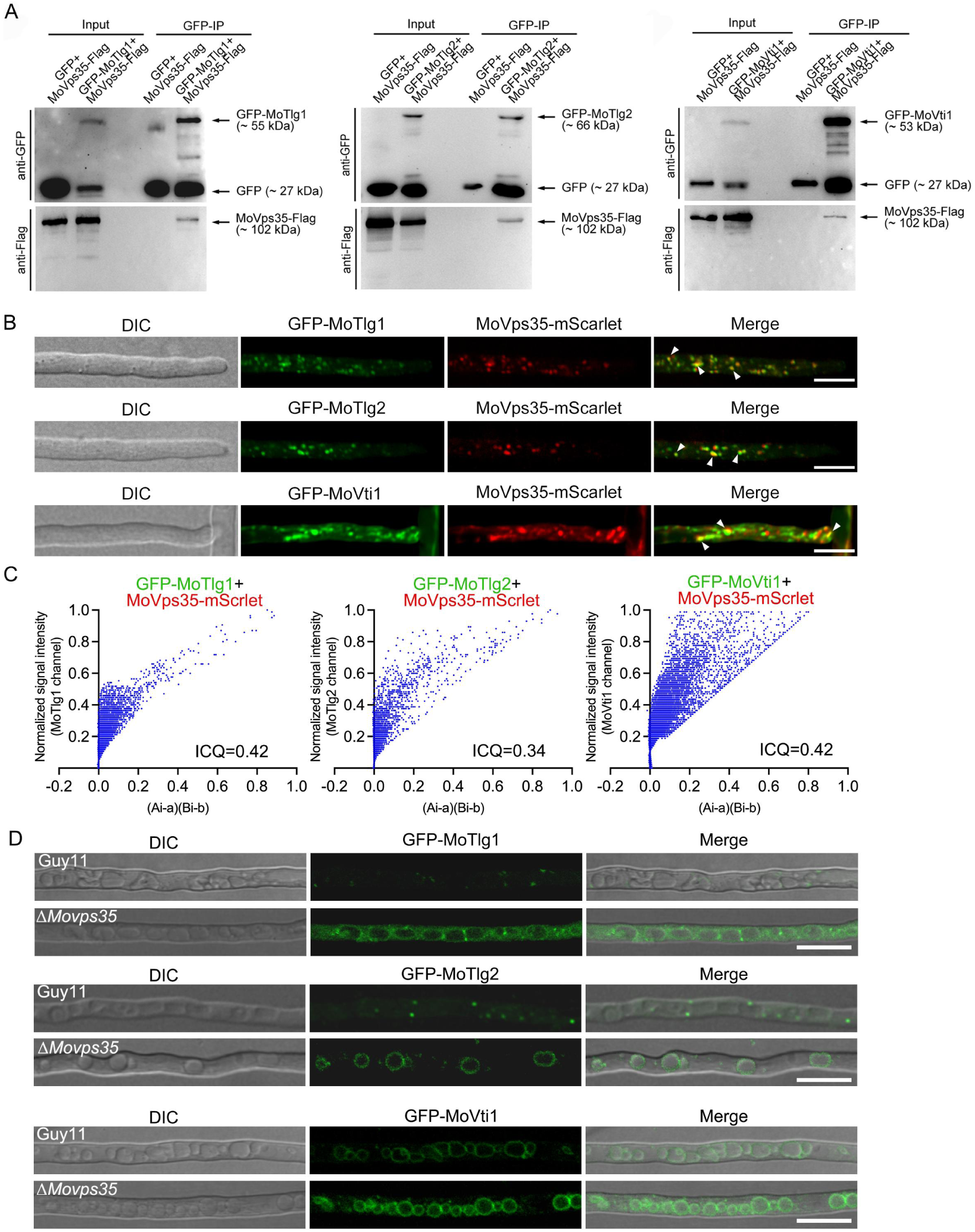
TGN-associated SNARE proteins interact with and are regulated by MoVps35. A. Coimmunoprecipitation showing positive interaction between MoVps35 and each of MoTlg1, MoTlg2 and MoVti1. Strains co-expressing the proteins were subjected to immunoprecipitation using GFP-trap beads. The immunoprecipitation signals (GFP-MoTlg1, GFP-MoTlg2, and GFP-MoVti1) alongside the Co-immunoprecipitation signal (MoVps35-Flag) were subsequently identified through immunoblot analysis employing antibodies specific to GFP and Flag, respectively. B. Co-localization assays of GFP-MoTlg1, GFP-MoTlg2, and GFP-MoVti1 with MoVps35-mScarlet, respectively, in the hyphae of *M. oryzae*. In the zoomed regions, the GFP-MoTlg1, GFP-MoTlg2, and GFP-MoVti1 partially co-localized (yellow) with MoVps35-mScarlet along the directions of the white arrows. Arrows indicate the colocalization of Green and mCherry fluorescence signals. C. A high degree of colocalization is indicated by a pixel cloud distribution of the plot. The plot graph illustrates the application of Li’s intensity correlation coefficient (ICQ), a metric employed for gauging the extent of colocalization. The terms ‘Ai’ and ‘a’ represent the current and average fluorescence intensities of the SNARE proteins channel, respectively, while ‘Bi’ and ‘b’ correspond to the respective values for the MoVps35 channel. The ICQ scale spans from −0.5, denoting exclusion, to 0.5, signifying complete colocalization. D. Localizations of GFP-MoTlg1, GFP-MoTlg2 and GFP-MoVti1 in the presence and absence of MoVps35. Deletion of *MoVPS35* disrupts the localizations of GFP-MoTlg1, GFP-MoTlg2 and GFP-MoVti1 compared to their localizations in the WT. The GFP-tagged MoTlg1, MoTlg2 and MoVti1 were observed to accumulate on the vacuolar membrane in Δ*Movps35* mutant strain, rather than the punctate structures observed in the WT. Scale bars = 10 µm.

### Scission of MoVps35-coated vesicles requires MoVps1 functions in *M. oryzae*

In yeast, Vps1 is a dynamin-like GTPase that plays a role in membrane scission events during vesicle formation and fission; and it has been reported to interact with Vps35 and other retromer components, suggesting a functional connection between Vps1-mediated membrane dynamics and the retromer-mediated cargo retrieval pathway [30]. However, Vps1 orthologs have not been functionally characterized in plant-pathogenic fungi. In our quest to unravel the intricate regulatory mechanisms of the retromer-SNARE pathway, we first validated the possible interaction between MoVps1 and MoVps35 in *M. oryzae*. Co-IP analysis confirmed the potential interaction between the two important proteins (Fig. 6A), suggesting a synergistic function of MoVps1 and MoVps35 in cellular transport of vesicles. Live cell imaging further revealed that MoVps35-mScarlet and MoVps1-GFP exhibit partial co-localization (Fig. 6B), and using Li’s correlation analysis, we found that both the cloud shape and data for MoVps1 (ICQ=0.36) are consistent with the partial colocalization of MoVps1 with MoVps35 (Fig. 6C). These results suggest a synergistic role of the two proteins during vesicle formation and transport.

**Fig 6.**
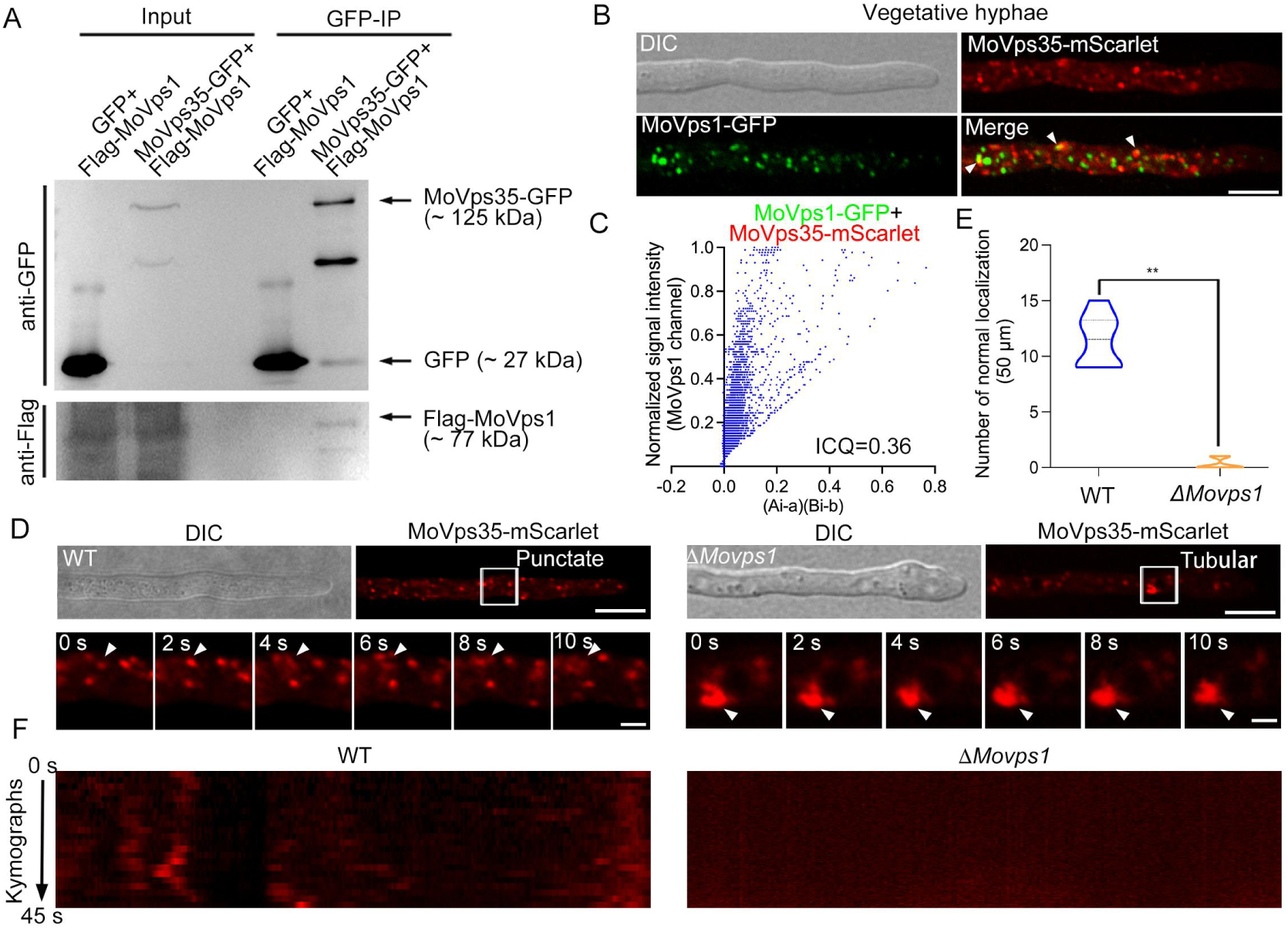
The dynamin-like GTPase MoVps1 regulates the fission of MoVps35-coated vesicles in *M. oryzae*. A.GFP-trap-based pull-down experiment indicating the physical interaction between MoVps35 and MoVps1. The immunoprecipitation signal (MoVps35-GFP) alongside the Co-immunoprecipitation signal (MoVps1-Flag) were subsequently identified through immunoblot analysis employing antibodies specific to GFP and Flag, respectively. B. Co-localization assay of MoVps35-mScarlet and MoVps1-GFP during vegetative growth. In the zoomed region, the MoVps1-GFP partially co-localized with MoVps35-mScarlet (yellow). Arrows indicate the colocalization of Green and mCherry fluorescence signal. C. A high degree of colocalization is indicated by a pixel cloud distribution of the plot. The plot graph illustrates the application of Li’s intensity correlation coefficient (ICQ), a metric employed for gauging the extent of colocalization. The terms ‘Ai’ and ‘a’ represent the current and average fluorescence intensities of the MoVps1 channel, respectively, while ‘Bi’ and ‘b’ correspond to the respective values for the MoVps35 channel. The ICQ scale spans from −0.5, denoting exclusion, to 0.5, signifying complete colocalization. D. Deletion of *MoVPS1* disrupts the normal localization of MoVps35-mScarlet. Signals categorized as “Punctate” exhibit punctate positioning, whereas those classified as “Tubular” display an extended tubular arrangement. Scale bars = 10 µm. E. Statistical analysis was performed to compare the number of punctate localizations of MoVps35-mScarlet between WT and Δ*Movps1* strains. F. Kymographs of “stream” time-lapse series of MoVps35-mScarlet in WT and Δ*Movps1* evidenced from S1 and S2 Videos, respectively.

To explore the function of the MoVps1-MoVps35 interaction, we examined the localization patterns of MoVps35-mScarlet in both the wild-type and Δ*Movps1* mutant strains. In the WT, MoVps35-mScarlet displayed a characteristic punctate localization pattern, indicative of its role in vesicle trafficking. In contrast, the retromer subunit exhibited an aberrant localization pattern, forming extended tubular structures in Δ*Movps1* mutant strain, signifying a disruption in the normal vesicle fission process (Fig. 6D-E). A kymograph further confirmed that the dynamics of MoVps35-mScarlet in Δ*Movps1* mutant are slower than in the wild type (Fig 6F). This demonstrates that MoVps1 might regulate the fission and proper distribution of MoVps35-coated vesicles. Taken together, our results demonstrate a direct physical interaction between MoVps1 and MoVps35 and that MoVps35 requires MoVps1 for its vesicle trafficking functions in *M. oryzae*. These results highlight the integral role of MoVps1 in the retromer-SNARE pathway in *M. oryzae*.

### MoVps1 is indispensable for the correct localizations of TGN-associated SNARE proteins in *M. oryzae*

To gain an insight into the regulatory relationship between MoVps1 and the TGN-associated SNARE proteins, we conducted co-localization studies to determine the spatial relationship between MoVps1 and the various TGN-associated SNARE proteins in the hyphae of *M. oryzae*. The results showed partial co-localization of MoVps1 with GFP-MoTlg2, GFP-MoVti1, GFP-MoTlg1, and GFP-MoSnc1, suggesting a potential role of MoVps1 in the trafficking or sorting of these proteins (Fig. 7A-B). To further explore the functional significance of MoVps1 in the localization of the TGN-associated SNARE proteins, we generated *MoVPS1* by genetic recombination (Fig. S6), and compared the localization patterns of the TGN-associated SNARE proteins in the wild-type and Δ*Movps1* mutant strains. In the WT strain, the SNARE proteins exhibited the typical punctate patterns consistent with their roles in vesicle trafficking. However, in the Δ*Movps1* mutant, there was a marked disruption in the normal localization of these proteins. GFP-MoTlg2 was mislocated to vacuoles, while GFP-MoVti1 and GFP-MoSnc1 showed abnormal accumulations often observed near the vacuolar regions; GFP-MoTlg1 was mis-localized to the cytosol and also accumulated around the vacuole, indicating a failure in their proper sorting and transportation (Fig. 7A and C). These findings demonstrate that MoVps1 is essential for the proper transportation and localization of the TGN-associated SNARE proteins in *M. oryzae*.

**Fig 7.**
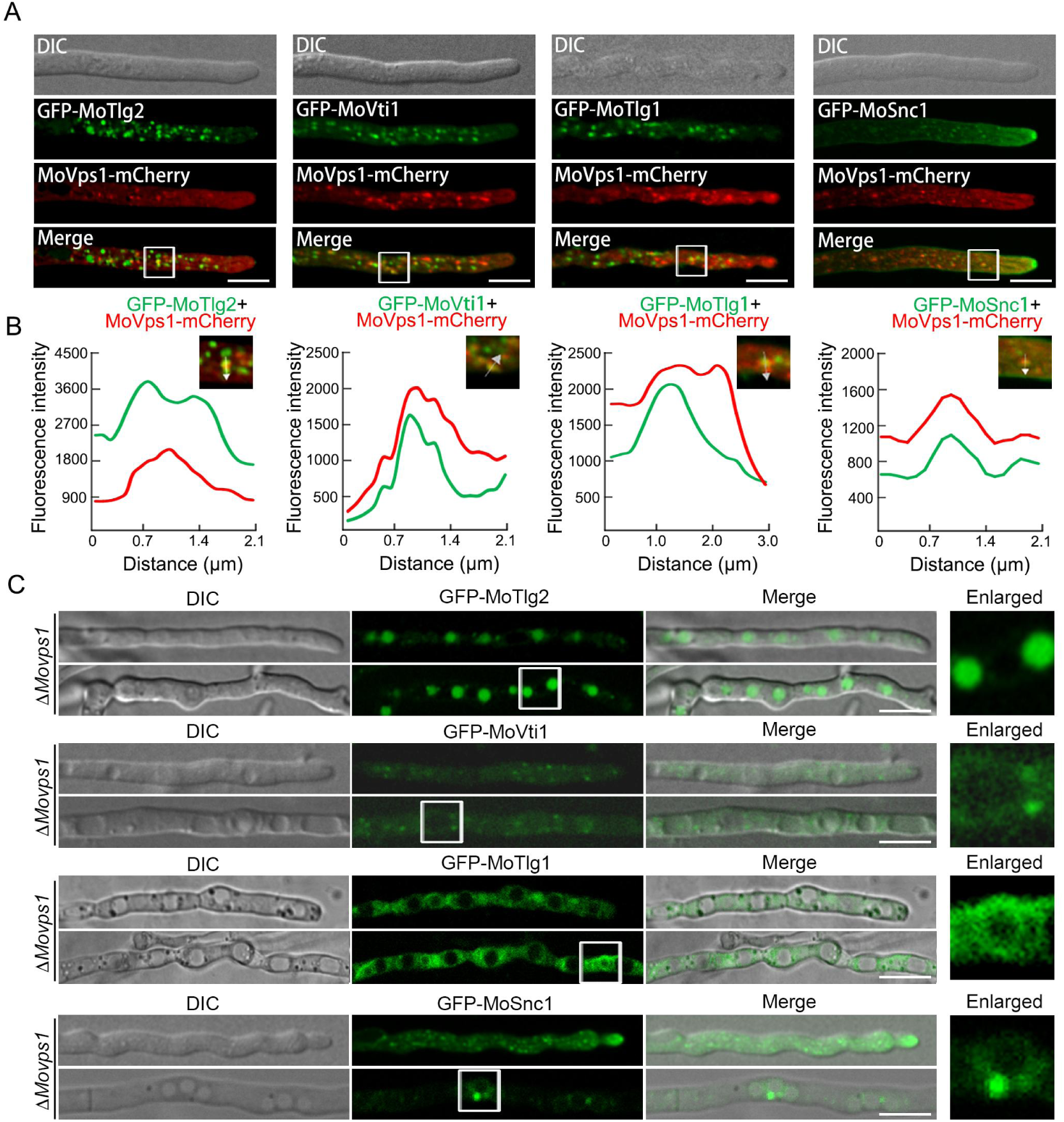
Loss of MoVps1 impairs the proper transportation of TGN-associated SNARE proteins. A. Co-localization of MoVps1-mCherry with GFP-MoTlg2, GFP-MoVti1, GFP-MoTlg1 and GFP-MoSnc1 in *M. oryzae* hyphae. B. Line scan graphs for the co-localization of MoVps1-mCherry with GFP-MoTlg2, GFP-MoVti1, GFP-MoTlg1 and GFP-MoSnc1. In the zoomed regions, GFP-MoTlg2, GFP-MoVti1, GFP-MoTlg1, and GFP-MoSnc1 partially co-localized with MoVps35-mScarlet (yellow). Each line scan graph was generated at the position indicated by the arrow in the zoomed region to show the relative fluorescence intensity of MoVps35-mScarlet with GFP-MoTlg1, GFP-MoTlg2, and GFP-MoVti1, respectively. C. Deletion of *MoVPS1* disrupts the localization of GFP-MoTlg2, GFP-MoVti1, GFP-MoTlg1, and GFP-MoSnc1 in fresh and old hyphae. The zoomed views show the details of the vacuole region. Scale bars = 10 µm.

### MoVps1 plays crucial roles in the development, asexual reproduction and pathogenicity of *M. oryzae*

To further investigate the biological functions of MoVps1 in the rice blast fungus, we analyzed the phenotypes of the Δ*Movps1* mutant and discovered that the mutant had a significantly reduced growth rate compared to the wild-type and complemented strains, indicating the importance of MoVps1 in the fungal vegetative development (Fig. 8A-B). More so, deletion of *MoVPS1* abolished asexual sporulation, as the wild-type and the complemented strains (cultured under same environmental conditions with the mutant) produced abundant and statistically similar amount of conidia while Δ*Movps1* mutant lacks such asexual structures (Fig. 8C). To assess the significance of the *MoVPS1* gene in the pathogenicity of *M. oryzae*, we performed hyphae-mediated infection assay by inoculating intact and injured barely leaves with mycelia harvested from the wild-type, Δ*Movps1* and Δ*Movps1* _Com strains. We similarly observed that the Δ*Movps1* mutant almost lost its virulence on both the intact and injured barley leaves compared to the wild-type and complemented strains. This highlights the pivotal role played by MoVps1 in the pathogenicity of *M. oryzae* (Fig. 8D).

**Fig 8.**
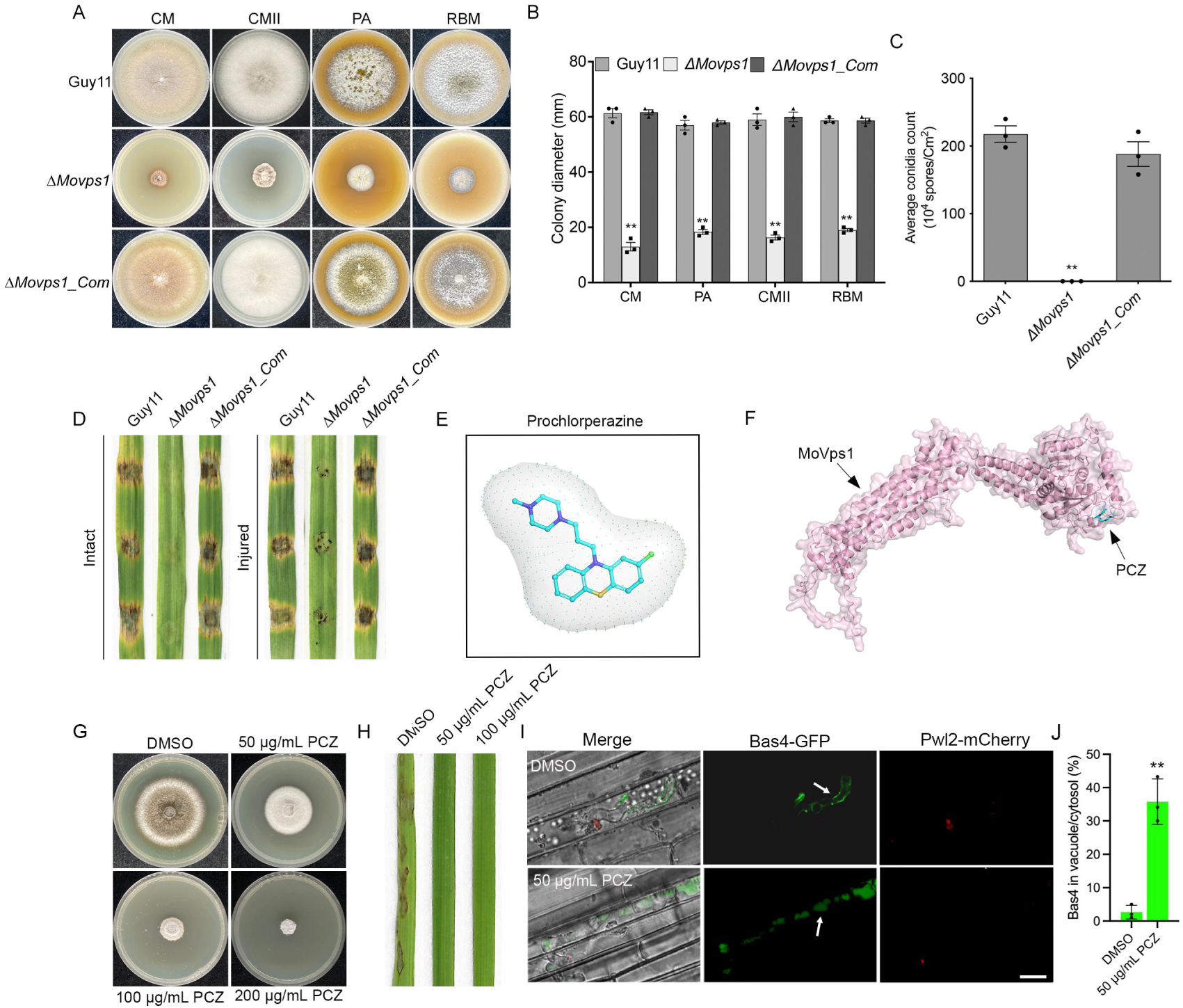
MoVps1 plays important role in the development, asexual reproduction and pathogenicity of *M. oryzae*. A. Analyses of the vegetative growths of the wild-type, Δ*Movps1* mutant and complemented strain Δ*Movps1_Com* on CM, CM II, PA and RBM media. B. Statistical analyses of the colony diameters of the indicated strains grown on CM, PA, CMII, and RBM for 10 days. C. Comparison of the number of conidia produced by the wild-type, Δ*Movps1* mutant and the complemented strain. The fungal strains were inoculated on RBM and incubated under 12 h/12 h light/dark condition for 3 days. D. Pathogenicity of Δ*Movps1* strain on intact and injured barley leaves compared to the wild-type and complemented strains. E. The structural formula of prochlorperazine (PCZ) as depicted by Pymol. F. The docking results between MoVps1 and PCZ by using AutoDocktools, which displayed a docking value of −6.94. G. Sensitivity of the wild-type strain to different concentrations of PCZ. The fungus was grown on CM media supplemented with the indicated concentrations of PCZ for 10 days. H. Analysis of the effect of PCZ on *M. oryzae* pathogenicity. The fungal spore suspensions (5×10^4^ spores/mL) were treated with DMSO, 50 μg/mL, and 100 μg/mL PCZ, respectively, and sprayed on 3-week-old susceptible CO39 rice seedlings. Disease lesions were observed and photographed at the 7th dpi. I. Relative localizations of Bas4-mCherry and Pwl2-mCherry following treatments with DMSO and 50 μg/mL PCZ. Arrows represent the localization of Bas4-GFP. Scale bars = 10 µm. J. Bar charts illustrating the percentage of Bas4 in vacuole in the DMSO and 50 μg/mL PCZ treatment. Error bars represent standard deviation from three replicates. Double asterisks denote significant differences at p< 0.01, based on t-test analysis.

Prochlorperazine (PCZ), a phenothiazine antipsychotic medication known to inhibit dynamin, disrupts vesicle scission from the plasma membrane, thereby impacting intracellular vesicle trafficking and cargo transport (Fig. 8E) [31, 32]. Given the established role of dynamin in vesicular transport, we hypothesized that PCZ might exert its effects on *M. oryzae* by interacting with proteins involved in vesicle transport, such as MoVps1.To further explore the molecular basis of this interaction, molecular docking techniques were used to investigate potential in silico interactions between PCZ and MoVps1. The analysis revealed the presence of a binding pocket on MoVps1 that could accommodate PCZ, suggesting a molecular mechanism by which PCZ may influence vesicle transport processes in *M. oryzae* (Fig. 8F). As such, we decided to check the sensitivity of the wild-type strain to PCZ. We observed that the wild-type strain exhibited a dose-dependent susceptibility to PCZ when cultured on CM medium supplemented with different concentrations of the drug, with the most notable inhibition of vegetative growth at the highest concentration used (Fig. 8G). Furthermore, we spray CO39 rice seedlings with wild-type Guy11 spores suspension which treated with 50 μg/mL and 100 μg/mL PCZ, respectively. The results revealed a near-complete loss of pathogenicity of the rice blast fungus (Fig. 8H). Further investigation into whether PCZ affects the secretion of effector proteins by *M. oryzae* revealed that, under 50 μg/mL PCZ treatment, Bas4-mCherry accumulated within the vacuoles of invasive hyphae while Pwl2-mCherry showed no difference in its localization after treatment with DMSO and 50 μg/mL PCZ, indicating that PCZ causes improper secretion and mis-localization of apoplastic effectors (Fig. 8I). These results suggest the potential of dynamin-like protein as a target for PCZ and support the repurposing of this phenothiazine derivative as a novel antifungal agent for rice blast disease management.

## Discussion

Our previous research has highlighted on the critical regulatory roles of the v-SNARE protein MoSnc1 in the transport and secretion of effectors within *M. oryzae,* which are crucial for its pathogenicity and adaptation to the host environment [13]. In this study, we broadened our perspective to encompass the broader dynamics of the SNARE complex, focusing in particular on the interplay between MoSnc1 and other TGN-associated SNARE proteins such as MoTlg1, MoTlg2 and MoVti1. Our results elucidate the crucial role of this SNARE complex in vesicle docking and fusion processes, which are essential for the secretion of apoplastic effectors. Furthermore, we have identified the dynamin-like GTPase MoVps1 and the retromer subunit MoVps35 as key regulators in the endomembrane trafficking system. Their absence leads to mis-localization of the TGN-associated SNARE proteins and has implications for pathogenicity (Fig. 9). Additionally, our research has uncovered the potential of phenothiazine as an antimicrobial agent to inhibit the pathogenic capability of *M. oryzae*. These results suggest a novel approach for the development of biopesticides targeting SNARE-mediated trafficking pathways.

**Fig 9.**
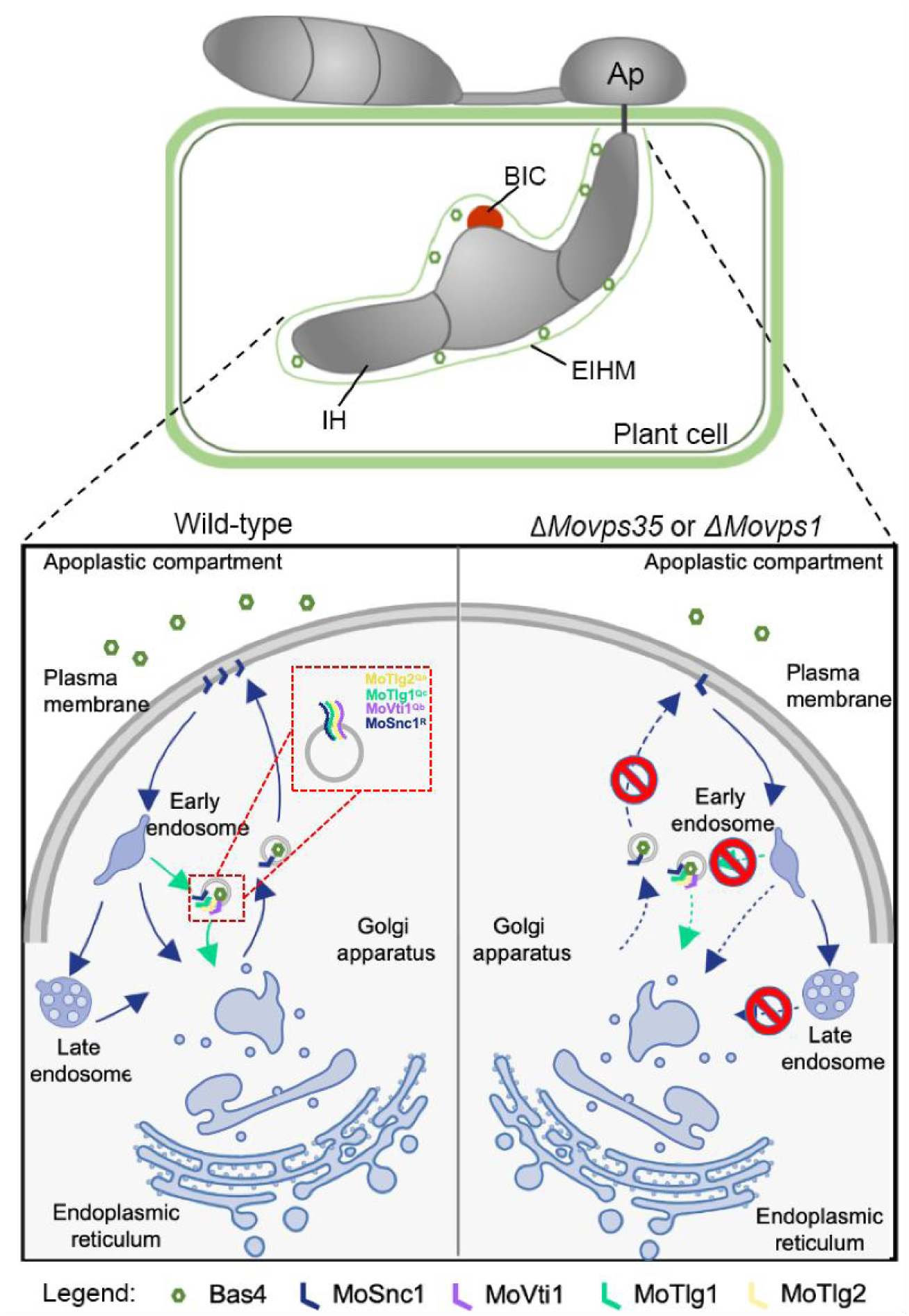
A proposed model for the synergistic roles of MoVps1 and MoVps35 during the trafficking of TGN-related SNARE proteins in *M. oryzae*. SNARE proteins play a pivotal role in vesicular trafficking process, particularly in the context of the trans-Golgi network. This process is crucial for regulating the secretion of apoplastic effectors in *M. oryzae*. In this model, MoVps1 and MoVps35 are posited to play key roles in the recycling and sorting of the TGN-associated SNARE proteins MoSnc1, MoTlg1, MoTlg2, and MoVti1, which are essential for vesicle fusion and cargo (such as effectors) delivery to the target membrane. Disrupting *MoVPS1* and *MoVPS35* results in detrimental effects on the functionality of the TGN-related SNARE proteins, leading to malfunctioning of the vesicular transport system within the rice blast fungus.

The vesicular transport system plays an essential role in the secretion of effectors in *M. oryzae* and facilitates the transport of these molecules to their respective compartments within the host organism, which is crucial for the ability of the pathogen to establish infection and exert its deleterious effects on rice plants [13, 14, 33]. The exocyst complex and SNARE proteins have been shown to function as docking subunits that tethering vesicles to the secretory sites [34, 35]. Notably, the endocytic protein MoEnd3 has been implicated in the secretion of the cytoplasmic effectors such as Avr-Pia and AvrPiz-t, but not for other apoplastic effectors like AvrPib and AvrPi9 [36]. Additionally, the Qc-SNARE protein MoSyn8 was found to be specifically required for the secretion of the cytoplasmic effectors Avr-Pia and AvrPiz-t, but not for apoplastic effectors [37]. Our research has identified a SNARE complex consisting of MoSnc1as well as MoTlg1, MoTlg2 and MoVti1, all of which are localized the TGN and collectively contribute to the precise targeting of apoplastic effectors. The specific reasons for this functional divergence between the SNARE proteins may be related to the unique protein-protein interactions and subcellular localization cues provided by each SNARE component provides. Thus, MoSyn8 could interact with a number of regulatory proteins or components of the vesicle-coating machinery that are specific for biogenesis and targeting of vesicles carrying cytoplasmic effectors. On the other hand, the MoSnc1/MoTlg1/MoTlg2/MoVti1 complex could be specifically recognized by Rab GTPases such as MoRab7 [13], which are known to regulate endolysosomal trafficking and effector secretion. Our study has shown that while the SNARE complex, consisting of MoSnc1, MoTlg1, MoTlg2, and MoVti1, is involved in the regulation of apoplastic effector secretion while both MoSnc1 and MoVti1 also exhibit regulatory roles in the secretion of cytoplasmic effectors (Fig. 4). This differential functionality is reflected in the diverse localization patterns of MoSnc1 and MoVti1 while perturbing the functions of MoVti1 was lethal to this phytopathogenic fungus (Figs. S2 and S4). In yeast, Snc1 and Vti1 are involved in the assembly of other SNARE complexes. For example, Snc1 can engage with other Q-SNAREs such as Sec9 and Sso1/Sso2 to mediate vesicle docking and fusion at the plasma membrane during exocytosis, and Vti1 involved with the formation of SNARE complexes with other proteins, such as Vam3, which mediate the fusion of endosomal vesicles with the vacuole [17], suggesting a parallel possibility of similar diversity within the SNARE complex machinery of *M. oryzae*.

The retromer is a protein complex that mediates the retrograde transport of cargo proteins from endosomes to the TGN or back to the plasma membrane [38]. Our previous study found 12 SNARE proteins to be highly enriched in MoVps35-GFP immunoaffinity purification assays, including MoSyn8 and MoSnc1 [13]. Further analysis revealed a potential interaction between a hypothetical TGN-associated SNARE complex, comprising MoSnc1, MoTlg1, MoTlg2 and MoVti1, and the retromer subunit MoVps35. Further investigations using Co-IP and live-cell imaging techniques confirmed these interactions (Fig. 5A-B), and disruption of *MoVPS35* resulted in mis-localization of the SNARE proteins into the vacuole membrane (Fig. 5C). This discovery is significant as it suggests a direct influence of the retromer complex on the assembly and trafficking of SNARE proteins, but the broader influence of the retromer complex on SNARE dynamics is still unclear. The retromer complex is composed of several subunits, including Vps35, Vps29, and Vps26, which together form a stable trimer. Disruption of the retromer complex can compromise the integrity and functionality of the Golgi apparatus, hindering the canonical effector secretion pathway [39]. Further research is required to uncovering the regulatory impact of the retromer complex on SNARE-mediated processes, which could potentially offer insights into cellular trafficking mechanisms and unveil therapeutic targets for the control of plant diseases.

Vps1 is a dynamin-like GTPase protein that plays a pivotal role in endomembrane trafficking, particularly in the late endosome-to-vacuole transport route [40]. It is involved in the fission of transport vesicles from donor compartments, which is a critical step in protein sorting and targeting within the cell [41]. Studies have shown that Vps1 and the retromer can collaborate to maintain vacuole homeostasis and ensure that proteins are efficiently recycled or targeted to their correct intracellular locations [41, 42]. Despite the well-documented importance of Vps1 in the context of the human pathogenic fungi, *Candida albicans*[43], its role in plant pathogenic fungi has remained unexplored. In *M. oryzae*, absence of *MoVPS1* leads to the emergence of elongated tubular structures and aggregate MoVps35-GFP (Fig. 6). This phenomenon may be attributed to the impaired function of MoVps1 in mediating the fission of retromer-coated vesicles from the endosomal membrane. Further analysis found that the absence of *MoVPS1* also causes mis-localization of TGN-associated SNARE proteins, resulting in a more complex and potentially disorganized subcellular distribution of the proteins (Fig. 7). In yeast, Vps1 can associate with the GARP (Golgi-associated retrograde protein) tethering complex, which includes Vps51 that facilitate endosome-to-Golgi transport [44]. These interactions highlight the complex regulatory network involving Vps1, which affects a range of downstream proteins that are crucial for vesicular trafficking. Therefore, the precise mechanism by which MoVps1 regulates the functions of TGN-associated SNARE proteins remains to be elucidated. Interestingly, we found in this study that MoVps1 serves as a potential therapeutic target of prochlorperazine, a small molecule that blocks endocytosis by inhibiting dynamin [45]. Treatment with PCZ was observed to cause mis-localization of apoplastic effectors to the vacuolar compartment (Fig. 8), implying that the mechanisms by which the retromer complex regulates the secretion of the two classes of effectors may depend on distinct upstream regulatory factors. It is conceivable that Vps1 functions in a pathway involving the retromer complex that has to do with secretion of apoplastic effectors. Treatment with PCZ may exert selective effects on the secretion of apoplastic effectors. This suggests that PCZ could modulate the activity or localization of Vps1 or its interacting partners among the retromer subunits, thereby disrupting the trafficking or secretion of apoplastic effectors without affecting the cytoplasmic ones. Further investigation is needed for clearer understanding of the precise molecular interactions and pathways.

In conclusion, our research has highlighted the critical roles of the dynamin-like GTPase MoVps1 and the retromer core component MoVps35 in regulating the trafficking of TGN-associated SNARE proteins. These proteins act in concert to ensure efficient secretion of apoplastic effectors, which is vital for the pathogenicity of *M. oryzae*. Our findings not only advance the understanding of effector secretion mechanisms but also open avenues for the development of novel biopesticides to control the rice blast disease. Future work could focus on the development of biopesticides by modulating the binding affinity or the molecular scaffold of prochlorperazine to efficiently target and disrupt the pathogenicity of *M. oryzae*.

## Materials and Methods

### Strains and growth conditions

The *M. oryzae* wild-type strains Guy11 and 70-15 were used to generate *MoTLG2* and *MoTLG1* deletion mutants, respectively. The genetically manipulated strains were cultured on complete medium (CM) plates containing 6 g yeast extract, 6 g casein hydrolysate, 10 g sucrose, and 20 g agar per liter at 28°C, while liquid CM (CM without agar) was used to obtain the fungal mycelia for DNA extraction and protoplast preparation. Additionally, rice bran media (RBM) containing 40 g rice bran and 20 g agar per liter at pH 6, and CMII (10 g D-glucose, 2 g peptone, 1 g yeast extract, 1 g casamino acids, 50 mL 20×nitrate salts, 1 mL trace elements, 1 mL vitamin solution and 15 g agar per liter, pH 6.5) were employed for assessing conidiation.

### Generation of gene deletion mutants and complementation

Targeted gene deletion using homologous recombination strategy was used to generate Δ*Motlg1* and Δ*Motlg2* mutants. First, we amplified approximately 1 kb DNA sequences upstream and downstream of the targeted open reading frames (ORFs) and cloned them into pCX62 vector harboring hygromycin resistance gene (for selection purposes), respectively. The resulting plasmids were utilized to generate the AH and HB fusion fragments, which were subsequently amplified and introduced into the protoplasts of the corresponding wild-type strains. Transformants were initially identified by PCR using OF/OR and UF/H853 primer pairs (listed in Table S1), and further verified by Southern blot analysis.

The GFP-MoTlg1 fusion vector was constructed by amplification of MoTlg1 coding sequence using the primers MoTlg1-GOF and MoTlg1-GOR, MoTlg1-PF and MoTlg1-PR primers were used to amplify the native promoter from the genomic DNA of the Guy11, and GFPF and GFPR primers were used to amplify the eGFP fragment from the pKNTG plasmid (listed in Table S1). The PCR products were cloned into pKNT vector using One Step Cloning Kit (Vazyme Biotech Co., Ltd) and verified by sequence analysis. The other fusion vector were constructed in similar protocol. The fusion vectors were sequenced to confirm the correct insertion and integrity of the inserted fragments, and then transformed into the protoplasts of their respective mutants by PEG-mediated transformation as described previously.

### Tet-off gene expression system

To develop the Tet-Off strain, the promoter and coding sequence of *MoVTI1* were cloned into a pFGL1252_TetGFP (Hyg) vector (Addgene ID 118992), which contains a Tet-off cassette that is responsive to tetracycline or doxycycline. The promoter of *MoVTI1* was PCR-amplified using the specific primers *MoVTI1* Tet-AF and *MoVTI1* Tet-AR. The PCR product was then cloned into the pFGL1252_TetGFP (Hyg) vector, which was pre-digested with *Xho* I and *Eco*R I, to generate pFGL1252_TetGFP (Hyg)-*MoVTI1*A construct. Subsequently, the approximately 1500-bp ORF of *MoVTI1* was amplified using the primers *MoVTI1* Tet-BF and *MoVTI1* Tet-BR, and this fragment was inserted into the pFGL1252_TetGFP (Hyg)-*MoVTI1*A vector, pre-digested with *Pst* I and *Hin*d III, yielding the final pFGL1252_TetGFP (Hyg)-*MoVTI1*AB construct. This construct was then transformed into wild-type protoplasts. The transformed protoplasts were initially screened and identified by PCR using specific primers (Table S1), followed by sequence analysis to confirm the correct insertion of the Tet-Off construct. Additionally, positive transformants were further validated by confocal microscopy to visualize the GFP fluorescence and to confirm the proper expression of the targeted gene under the control of the Tet-Off promoter.

To validate the functionality of the Tet-Off system, the transformants were treated with tetracycline or doxycycline and analyzed for changes in the expression of the gene of interest, as indicated by loss or reduction in fluorescence. This phenotypic analysis confirmed the successful establishment of the Tet-Off strain, which allows for the regulated expression of the target gene in response to the presence or absence of tetracycline or doxycycline.

### Assays for vegetative growth and conidiation

For assessment of vegetative growth, both the wild-type and genetically modified strains, including the mutants and complemented lines, were cultivated on four distinct media: CM, prune agar (PA, per L: 40 mL prune juice, 5 g lactose, 5 g sucrose, 1 g yeast extract and 20 g agar, pH 6.5), CM II, and RBM. The cultures were incubated at 28°C under 12 h/12 h light-dark cycles. The diameters of the fungal colonies were measured 10 days post incubation.

For conidiation assay, the strains under study were inoculated on RBM and maintained under continuous dark for a 7-day period. Subsequently, the hyphae were gently scraped using sterilized glass slides to dislodge the conidia, and the plates were then exposed to 12 h light-dark cycles to promote conidial release and collection. The conidia were harvested by washing the surfaces of the media with ddH_2_O, and the resulting suspension was filtered through a single layer of lens cleaning paper to remove any media debris. The volume of the conidial suspension was then adjusted to 2 mL for uniformity in subsequent analyses.

To facilitate the observation of conidiophore, the hyphae were scratched onto the surface of the media, and blocks of the media were transferred onto microscope slides for microscopic examination. These slides were incubated at 28°C under light conditions to allow for the development of conidiophores. The formation and morphology of conidiophores were assessed at 24 hours post-incubation (hpi) using a light microscope.

### Plant infection and penetration assays

For hyphae-mediated infection experiments, the individual strains were initially cultured in liquid CM for 3 days under constant agitation at 120 rpm, 28℃. Following this, the mycelia were harvested by filtration and rinsed thoroughly with sterile ddH_2_O to remove any residual media. Then, the media-free mycelia were used as propagules to inoculate intact and injured barley leaves. The inoculated barely were first incubated in a dark and humid condition for 24 h before being transferred to a growth chamber with a photoperiod of 12 h light-dark cycles for 5 days.

For conidia-mediated infection assays, spore suspensions were prepared at a concentration of 5×10^4^ spores per mL, supplemented with 0.02% (v/v) Tween-20. The conidia suspensions were used to inoculate the 7 days old intact and injured barely leaves, as well as to spray on 3 weeks old rice seedlings (cv. CO39). After inoculation, the seedlings were kept in a dark and humid environment for the initial 24-hour period, after which they were moved to a growth chamber with alternating 12 h light-dark cycles photoperiod for 5 days.

For penetration assays, spore suspensions were prepared in a similar manner to those used for the conidia-mediated infection assays on rice seedlings, except for the omission of Tween-20 supplementation. The spore suspensions were infiltrated into the sheaths of 4 weeks old rice and maintained in a dark and humid condition for 24 h. Development of invasive hyphae was observed under a light microscope.

### Co-localization assay

To validate the subcellular localization of TGN-associated SNAREs in *M. oryzae*, MoKex2-mCherry, GFP-MoTlg1, GFP-MoTlg2, and GFP-MoVti1 fusion proteins were generated. For GFP-MoSnc1, the vector constructed in our previous study [13] was used. Pair of vectors (GFP-MoTlg1/TGN-mCherry, GFP-MoTlg2/TGN-mCherry and GFP-MoVti1/TGN-mCherry) were co-transformed into Guy11 protoplasts, respectively. Transformants were initially identified through PCR screening, which was followed by examination of GFP signals by confocal microscopy to confirm the successful integration and expression of the constructs. Furthermore, to ascertain the co-localization patterns of the TGN-associated SNAREs with other key components of the trafficking pathway (such as MoVps35 and MoVps1), similar strategy was adopted. The strains intended for co-localization studies were generated and analyzed as outlined in the initial description.

To estimate colocalization, we used Li’s intensity correlation analysis, using the ImageJ JACoP plugin[46]. This analysis assesses the covariance between pixel intensities in two different channels, yielding an intensity correlation coefficient (ICQ) theoretically ranging from −0.5 (exclusion) to 0.5 (complete colocalization), with 0 value standing for no correlation (random staining). Intensity correlation plots created by Graphprime 10 can be more informative than ICQs particularly in these cases where the contribution of cytoplasmic and intranuclear pixels to the ICQ is substantial, because the shape of the cloud in the plots reveals whether ‘colocalizing’ values are due to co-variance of low intensity pixels (probably representing cytoplasmic background) or of high intensity pixels (the ‘true’ signals of the markers).

### Yeast two-hybrid assay

To investigate the potential interactions among MoSnc1, MoTlg1, MoTlg2, and MoVti1, the cDNA sequences of the genes encoding these proteins were cloned individually into the bait vector pGBKT7 and the prey vector pGADT7, allowing for the assessment of pairwise interactions. The interaction of pGBKT7-p53 and pGADT7-T was used as positive control while that of pGBKT7-Lam and pGADT7-T was used as negative control. The resulting bait and prey vectors were confirmed by sequencing, and co-transformed into the yeast strain AH109. All transformed cells were cultured on SD−Trp−Leu and SD−Trp−Leu−His−Ade plates, incubated at 30°C for 3 days. After selecting colonies, the cells were adjusted to 1×10^6^ cells/mL, after which droplets were placed on SD−Trp−Leu and SD−Trp−Leu−His−Ade plates containing 20 mg/mL X-α-Gal and further incubated at 30°C for 3 days.

### Co-immunoprecipitation assa**y**

Total proteins were extracted from the strains expressing GFP-MoSnc1 and GFP. These proteins were then incubated with 30 μL of anti-GFP magnetic agarose beads (Smart-Life Sciences, China) for 4 hours at 4°C to allow for the binding of GFP-tagged proteins. The beads were then collected and washed three times with 500 μL of cold washing buffer (50 mM Tris, 0.15 M NaCl, pH 7.4) to remove any unbound proteins or contaminants. A magnetic frame was used to further wash the beads three times with 500 μL cold washing buffer (50 mM Tris, 0.15 M NaCl, and pH 7.4). The beads were then resuspended in 100 μL washing buffer containing SDS-loading buffer. The proteins bound to the beads were then eluted by heating at 100°C for 15 min. The eluted protein samples were analyzed by immunoblot detection using anti-GFP antibodies (Abmart, China), followed by mass spectrometry analysis (Fujian Agriculture and Forestry University Analysis and Testing Center, China).

### Microscopy examinations

Nikon Ci-S fluorescence microscope and Nikon CUS-W1 spinning-disk confocal microscope (Japan), were utilized for the observation of GFP and mCherry fluorescence. The emission and excitation wavelengths used were 488 nm and 561 nm, respectively.

## Author contributions

L.L, D.L, Z.W., and W.Z. designed and sourced funding for the research, W.Z. secured the funding, L.L, Q.W., X.H., S.W., Q.G., Y.L., Y.A., and J.C. performed the experiments. L.L, W.Z., and Q.W. analyzed the data. L.L drafted the manuscript. L.L, D.L., W.Z., and X.H. revised the manuscript. All authors contributed to the final manuscript.

## Acknowledgments

We are grateful to the members of W.Z. laboratory for their insightful discussions and the technical support from Instrumental Analysis Center of Fujian Agriculture and Forestry University. No conflict of interest is declared.

