## Supplementary material for "Recycling of trans-Golgi SNAREs is essential for apoplastic effector secretion and effective pathogenicity of *Magnaporthe oryzae*": Table S1

**Table S1. The primers used in this study**

| Primers | Sequence（5'-3'） | Application |
| --- | --- | --- |
| MoTlg1-AF | TTGCGAGCCGTTAGTGTC | *MoTLG1* deletion |
| MoTlg1-AR | TTGACCTCCACTAGCTCCAGCCAAGCCGTTGAGGTGGGTCCGATT |  |
| MoTlg1-BF | GAATAGAGTAGATGCCGACCGCGGGTTTCCCAGTTGGAGTGATGAT |  |
| MoTlg1-BR | TCCGAGGAAACGACAAGA |  |
| YG/F | GATGTAGGAGGGCGTGGATATGTCCT |  |
| HY/R | GTATTGACCGATTCCTTGCGGTCCGAA |  |
| HYG/F | GGCTTGGCTGGAGCTAGTGGAGGTCAA |  |
| HYG/R | AACCCGCGGTCGGCATCTACTCTATTC |  |
| MoTlg1-OF | CGTCACCACCACAGATTTAC | Δ*Motlg1* mutant screen |
| MoTlg1-OR | AAACTACTCCACCGATCCTC |  |
| MoTlg1-UF | CAACCAACTGCGGAGATA |  |
| H853 | GACAGACGTCGCGGTGAGTT |  |
| MoTlg1-PF | AGGGAACAAAAGCTGGGTACTCGCGAGTCCTGGTCCAGCG | GFP-MoTlg1 generation |
| MoTlg1-PR | TCCTCGCCCTTGCTCACCATCCTGCGAATAGTGTTGCTCC |  |
| GFP-F | ATGGTGAGCAAGGGCGAGGA |  |
| GFP-R | CTTGTAGCTCGTCCATGC |  |
| MoTlg1-GOF | GCATGGACGAGCTGTACAAGATGATGTCCTCCACTAACGA |  |
| MoTlg1-GOR | CAGTAACGTTAAGTGGATCCTTGCGGTATTTTCCAAATTA |  |
| MoTlg2-AF | GAACAAAAGCTGGGTCTAACGGAACGCCAGAAT | *MoTLG2* deletion |
| MoTlg2-AR | CAGCGGCGCGCCGAACGAGTAGGTAGCGGACAGA |  |
| MoTlg2-BF | ACCGGGCCGGCCGGACACATCTCAACGGGTCA |  |
| MoTlg2-BR | GGTGGCGGCCGCTCTTCTACGCCTCAACAAGC |  |
| MoTlg2-OF | CTCCTACCGCCAGTCGT | Δ*Motlg2* mutant screen |
| MoTlg2-OR | TCTATGATGCCCTGTGCTAT |  |
| MoTlg2-UF | CTGGCTTGGCTGGGACA |  |
| MoTlg2-PF | GGGTACCGGGCCCCCCCTCGAGAGCCTCTTATCACAACTG | GFP-MoTlg2 generation |
| MoTlg2-PR | AGTTCCTCGCCCTTGCCCATGGCGGACCGCGAGTAG |  |
| MoTlg2-GOF | GCATGGATGAACTCTACAAGATGTGGCGAGATCGCACCAACCT |  |
| MoTlg2-GOR | CCCCCGGGCTGCAGGAATTCTCACCCACCATCGCCCGAGTCT |  |
| MoVti1-tetAF | ATTATTATGGAGAAACTCGAGCAAGACTTACACAGAACTGG | *MoVTI1* deletion by Tet-off system |
| MoVti1-tetAR | ATCCAGGCGGGCCATGAATTCGGCGAAGCGGGTATTTGGTG |  |
| MoVti1-tetBF | AGTTCTAGAGTCGACCTGCAGATGTCCAACCCCCTCGACGC |  |
| MoVti1-tetBR | ACGACGGCCAGTGCCAAGCTTCAAACACCAGCAGACCTTCA |  |
| MoVti1-tetF1 | GCAACTAGCTAATGCACTAG | *MoVTI1* Tet-off mutant screen |
| MoVti1-tetR1 | ACCTCGTTCAGCAGCTCCAA |  |
| MoVti1-tetF2 | ATGGTGAGCAAGGGCGAG |  |
| MoVti1-tetR2 | TTTTATGCTGTCGTCACGTT |  |
| MoVti1-PF | AGGGAACAAAAGCTGGGTACCAAGACTTACACAGAACTGG | GFP-MoVti1 generation |
| MoVti1-PR | TCCTCGCCCTTGCTCACCATGGCGAAGCGGGTATTTGGTG |  |
| MoVti1-GOF | GCATGGACGAGCTGTACAAGATGTCCAACCCCCTCGACGC |  |
| MoVti1-GOR | CAGTAACGTTAAGTGGATCCTTTTATGCTGTCGTCACGTT |  |
| MoVps1-AF | GAACAAAAGCTGGGTTTAGGATGCCGTTGAGG | *MoVPS1* deletion |
| MoVps1-AR | CAGCGGCGCGCCGAAGCACCGATGAATGCTTAT |  |
| MoVps1-BF | ACCGGGCCGGCCGGATTGAGAATCGGAACTGATAAAC |  |
| MoVps1-BR | GGTGGCGGCCGCTCTAGCCCTGTAATAACCTGTAAGC |  |
| MoVps1-OF | TCGTGGGCAGTCAATCAA | Δ*Movps1* mutant screen |
| MoVps1-OR | TTCAGACCGTCCGAGTTT |  |
| MoVps1-UF | AGTCGGTAATTCTCGGTTCT |  |
| MoVps1-GF | AGGGAACAAAAGCTGGGTACCCGCACGACGGCACAAAG | MoVps1-GFP generation |
| MoVps1-GR | GCCGCCGCCGCCGCCAAGCTTCTGCACCTGTCCGACAATTT |  |
| MoVps1-mF | TATAGGGCGAATTGGGTACC CGCACGACGGCACAAAG | MoVps1-mCherry generation |
| MoVps1-mR | CTTGCTCACCATAAGCTTGCCGCCGCCGCCGCCGCCGCCCTGCACCTGTCCGACAATTT |  |
| MoKex2-mCF | TATAGGGCGAATTGGGTACCCGAAACAAGGAAGACAAAGA | TGN-mCherry generation |
| MoKex2-mCR | CCCTTGCTCACCATAAGCTTTCGTGAAGTCATAGGCC |  |
| MoSnc1-ADF | GACGTACCAGATTACGCTCATATGCCCGAAGACGCTC | For generation of pAD-MoSnc1 and pBD-MoSnc1 construct |
| MoSnc1-ADR | TATCGATGCCCACCCGGGTGGAATTAGTTGCCCTTGAAGTGGA |  |
| MoSnc1-BDF | CTGATCTCAGAGGAGGACCTGCATATGCCCGAAGACGCTC |  |
| MoSnc1-BDR | CGCTGCAGGTCGACGGATCCCCGGGAATTAGTTGCCCTTGAAGTGGA |  |
| MoTlg1-ADF | GACGTACCAGATTACGCTCATATGATGTCCTCCACTAACG | For generation of pAD-MoTlg1 and pBD-MoTlg1 construct |
| MoTlg1-ADR | TATCGATGCCCACCCGGGTGGAATCATAACACAAGCAACAAGA |  |
| MoTlg1-BDF | CTGATCTCAGAGGAGGACCTGCATATGATGTCCTCCACTAACG |  |
| MoTlg1-BDR | CGCTGCAGGTCGACGGATCCCCGGGAATCATAACACAAGCAACAAGA |  |
| MoTlg2-ADF | GACGTACCAGATTACGCTCATATGTGGCGAGATCGCACC | For generation of pAD-MoTlg2 and pBD-MoTlg2 construct |
| MoTlg2-ADR | TATCGATGCCCACCCGGGTGGAATCACCCACCATCGCCC |  |
| MoTlg2-BDF | CTGATCTCAGAGGAGGACCTGCATATGTGGCGAGATCGCACC |  |
| MoTlg2-BDR | CGCTGCAGGTCGACGGATCCCCGGGAATCACCCACCATCGCCC |  |
| MoVti1-ADF | GACGTACCAGATTACGCTCATATGTCCAACCCCCTCG | For generation of pAD-MoVti1 and pBD-MoVti1 construct |
| MoVti1-ADR | TATCGATGCCCACCCGGGTGGAACTACCTGAACTTGCTAACA |  |
| MoVti1-BDF | CTGATCTCAGAGGAGGACCTGCATATGTCCAACCCCCTCG |  |
| MoVti1-BDR | CGCTGCAGGTCGACGGATCCCCGGGAACTACCTGAACTTGCTAACA |  |
| MoVps35-PmSF | GGGAACAAAAGCTGGGTACCTGGCAGATGCTTCTCACTCAG | For generation of MoVps35-mScarlet construct |
| MoVps35-PmSR | CCCTTGCTCACCATCTCGAGCTTGGGATCCAACACAATTCC |  |
| MoVps1-flagF | ATGACGACAAAGGTGGTAAGATGGCAGCCCAGACTTC | For generation of pFlag-MoVps1 construct |
| MoVps1-flagR | TCCTCGCCCTTGCTCACGAATTACTGCACCTGTCCGAC |  |
| MoVps1-hisF | CCATGGCTGATATCGGATCCATGGCAGCCCAGACTTCTCT | For generation of pHis-MoVps1 construct |
| MoVps1-hisR | TCGAGTGCGGCCGCAAGCTTTTACTGCACCTGTCCGACA |  |
| MoVps1-mbpF | ATCGAGGGAAGGATTTCAGAATTCATGGCAGCCCAGACTTCTCT | For generation of pMbp-MoVps1 construct |
| MoVps1-mbpR | GCCAGTGCCAAGCTTGCCTGCAGTTACTGCACCTGTCCGACA |  |
