## Supplemental information for "Recycling of trans-Golgi SNAREs is essential for apoplastic effector secretion and effective pathogenicity of *Magnaporthe oryzae*"

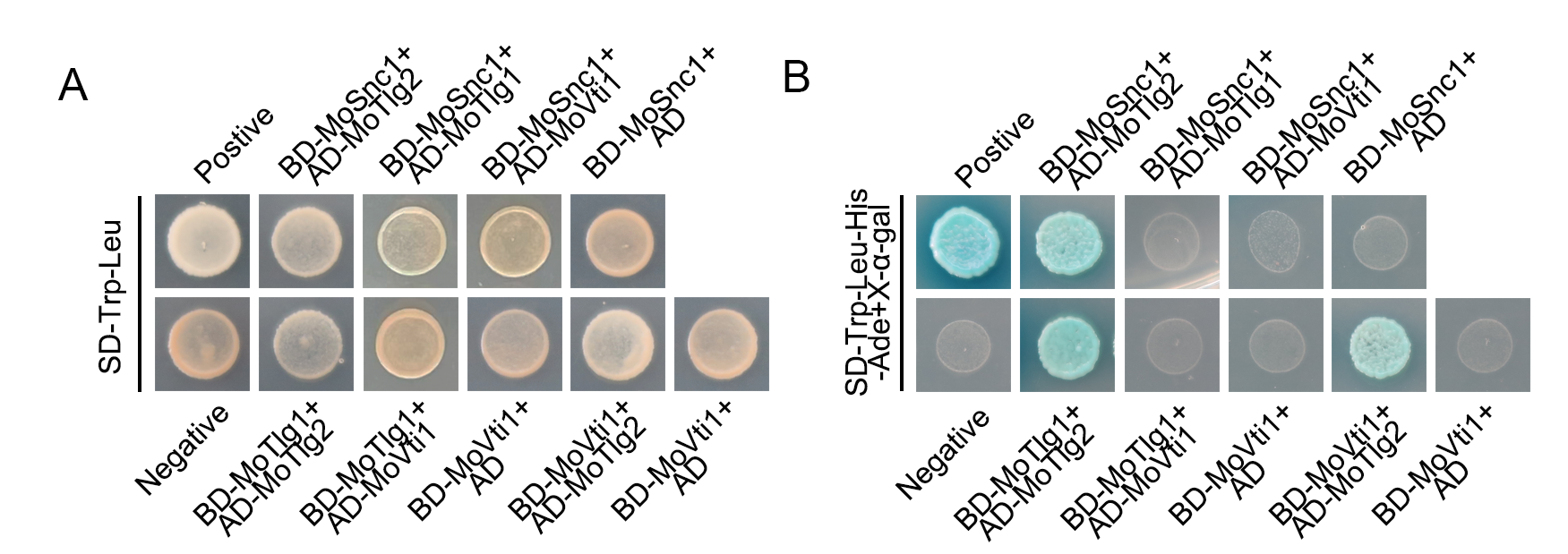

**Fig S1. Yeast-two-hybrid (Y2H) assays demonstrating the interaction patterns among MoSnc1, MoTlg1, MoTlg2, and MoVti1 proteins.**

A. Yeast-two-hybrid assay of the growth yeast strains harboring SNAREs-BD-paired SNAREs-AD on synthetic defined (SD) minimal yeast media plates supplemented with Trp-Leu as a measure of interaction between the paired subunits. Panel B shows the growth of yeast transformants expressing the SNAREs-BD-paired SNAREs-AD on SD supplemented with Trp-Leu-His-Ade + X-α-gal.

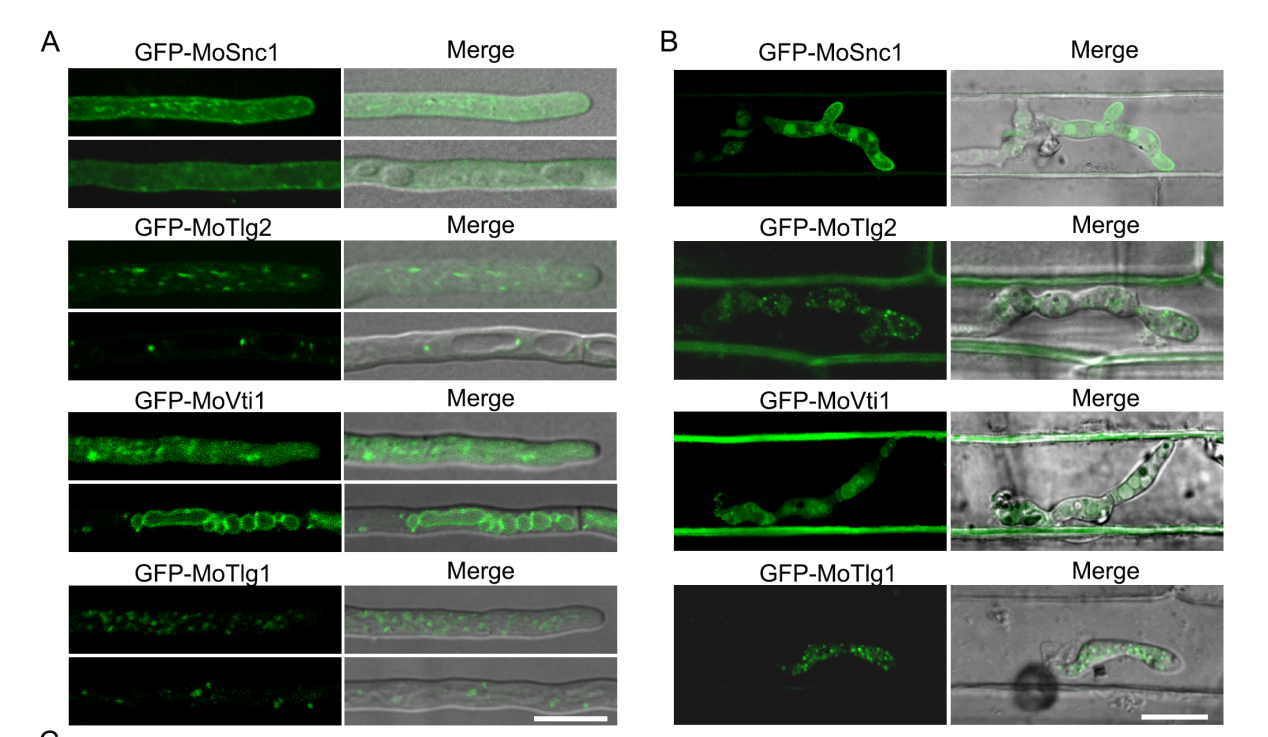

**Fig S2. Subcellular localization of GFP-MoSnc1, GFP-MoTlg2, GFP-MoVti1, and GFP-MoTlg1 in *M. oryzae***

A. Detailed confocal microscopy showing the subcellular localization of GFP-tagged SNARE proteins, including GFP-MoSnc1, GFP-MoTlg2, GFP-MoVti1, and GFP-MoTlg1, within distinct regions of the fungal hyphae. B. Subcellular localization of GFP-MoSnc1, GFP-MoTlg2, GFP-MoVti1, and GFP-MoTlg1 during the early infection stage of *M. oryzae*. Bar=10 μm.

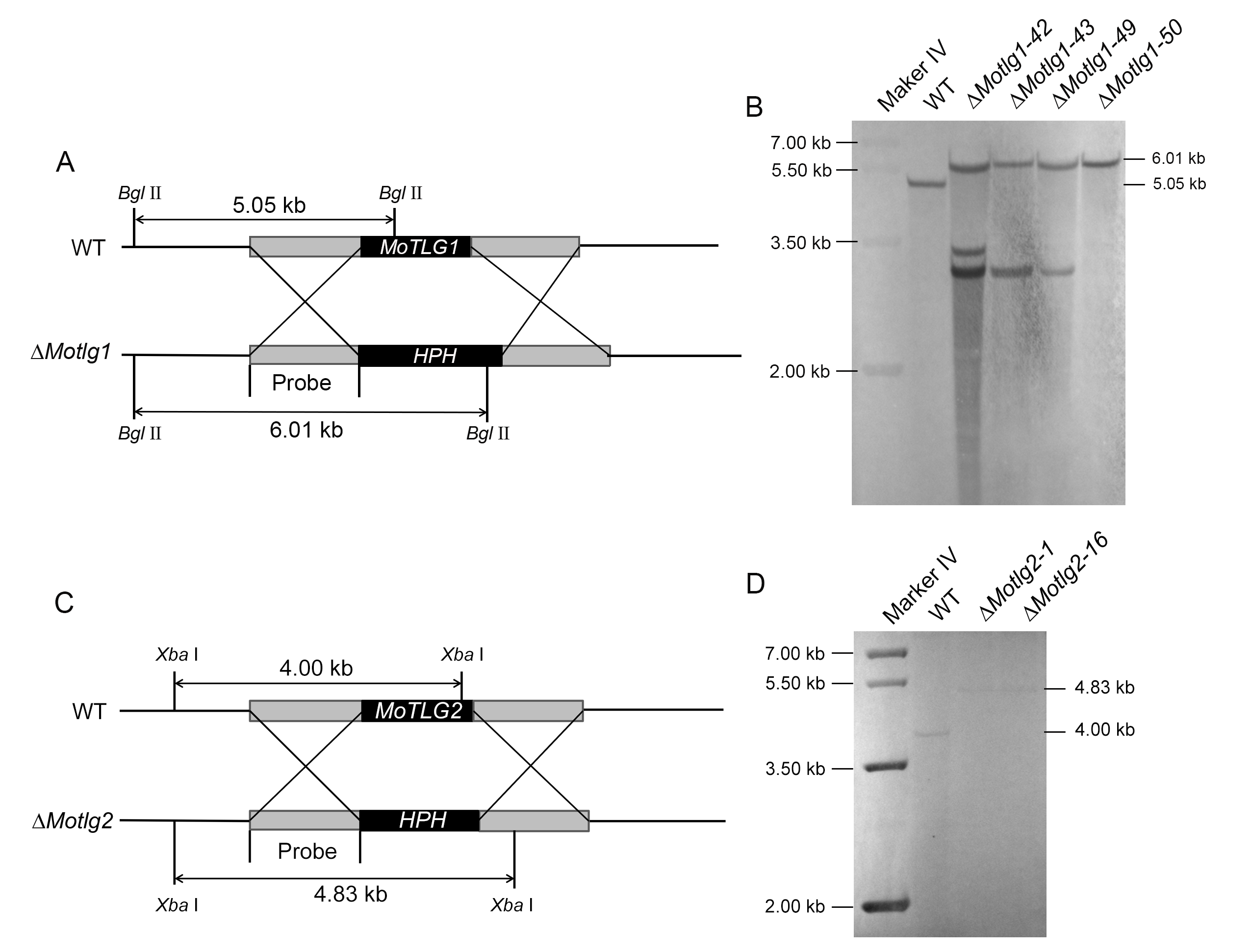

**Fig S3. Targeted gene deletion of *MoTLG1* and *MoTLG2* in *M. oryzae***

A. Schematic representation of the targeted gene disruption strategy used to generate Δ*Motlg1* strains by homologous recombination. B. Southern blot result confirming the successful replacement of *MoTLG1* with hygromycin resistance gene in the Δ*Motlg1* strains by single insertion, and a successful reintroduction of *MoTLG1* gene into the Δ*Motlg1* strains. C. Schematic representation of targeted gene disruption strategy used to generate Δ*Motlg2* strains by homologous recombination. D. Southern blot result confirming the successful replacement of *MoTLG2* with hygromycin resistance gene in the Δ*Motlg2* strains by single insertion and a successful reintroduction of *MoTLG2* gene into the Δ*Motlg2* strains.

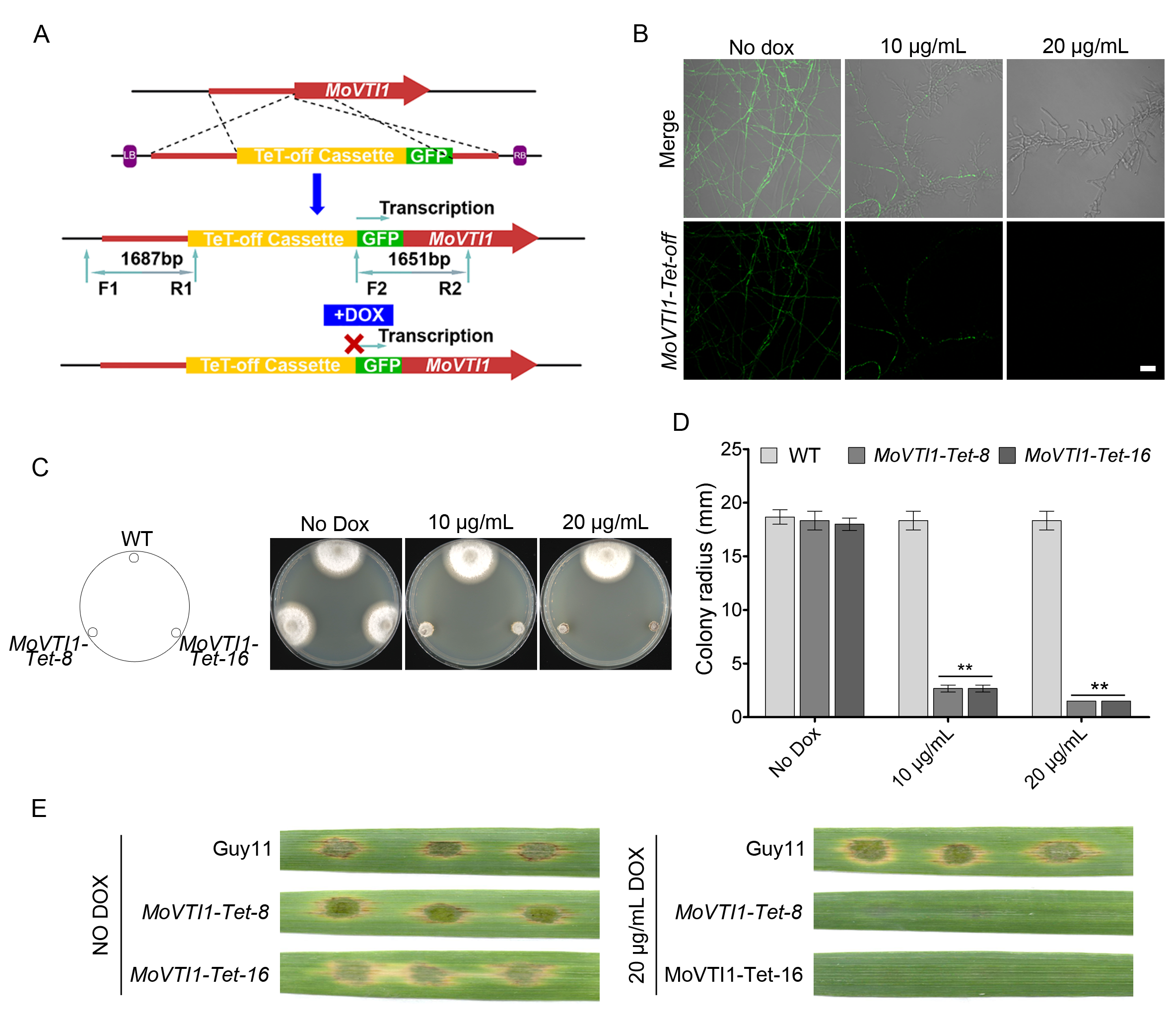

**Fig S4. Generation and phenotypic analyses of *MoVTI1-Tet* mutants**

A. Schematic diagram representing the gene silencing of *MoVTI1.* Primers F1, R1, F2 and R2 were used to screen and identify the correct transformants. B. Confocal examination to detect the signal of *MoVTI1-Tet* mutants in the presence or absence of difference concentrations of doxycycline (DOX). The GFP fluorescence was diminished progressively with the administration of escalating concentrations of DOX. C. Confirmation of successful silencing of *MoVTI1* by testing the growth of the mutants on CM media with or without doxycycline (DOX). D. Analysis of the average growth rates of the various mutant strains and the wild-type under different concentrations of DOX. E. Testing the pathogenicity of the *MoVTI1-Tet* mutant in the presence or absence of DOX. Conidia suspensions from the indicated strains were inoculated on barley leaves under treatment with 0 μg/mL and 20 μg/mL doxycycline. Photos were taken 7 days after inoculation.

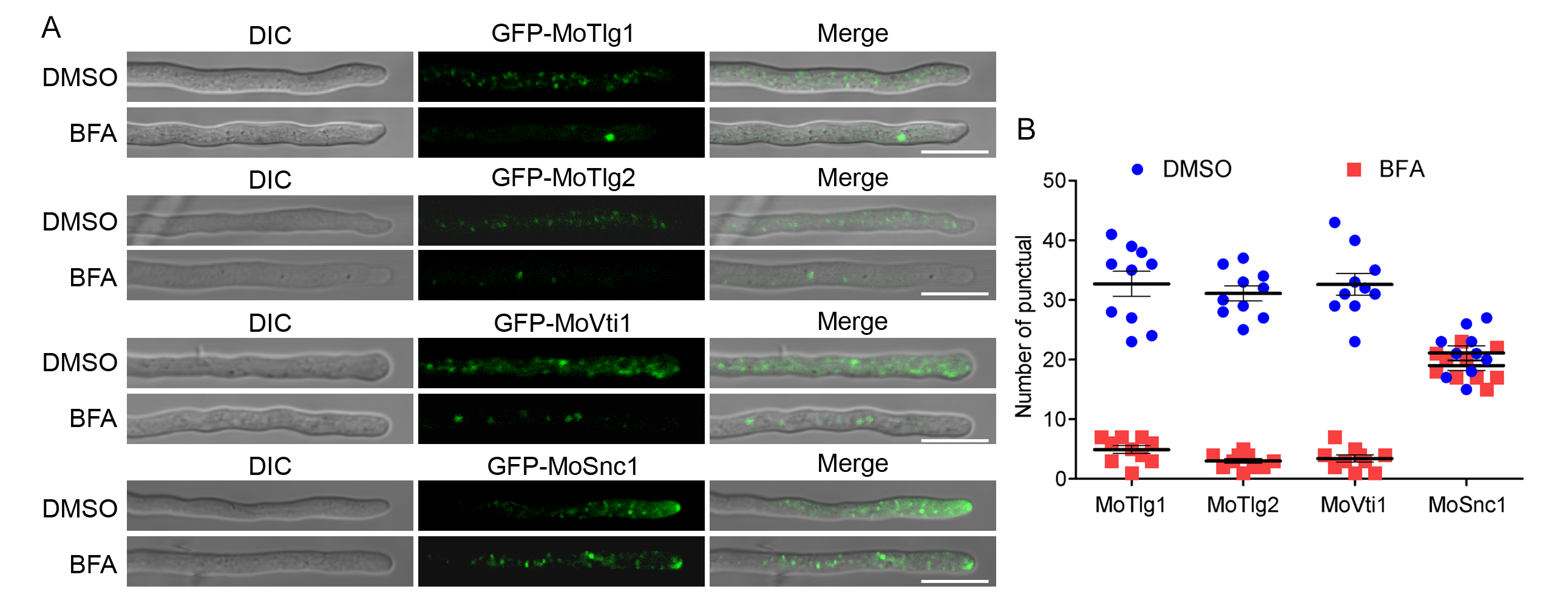

**Fig S5. Brefeldin A (BFA) inhibitor treatment disrupted the localization of TGN-associated SNARE proteins**

A. Illustration of the effect of Brefeldin A (BFA), a vesicular transport inhibitor, on the normal trafficking patterns of the GFP-tagged TGN-associated SNARE proteins GFP-MoTlg1, GFP-MoTlg2, GFP-MoVti1 and GFP-MoSnc1. The application of BFA notably disrupted the typical localization and distribution of these proteins within the hyphae. B. Quantitative analysis of the number of punctate structures (indicative of protein localization sites) observed in individual fluorescent strains following treatment with dimethyl sulfoxide (DMSO, as a control) and BFA. Bar=10 μm.

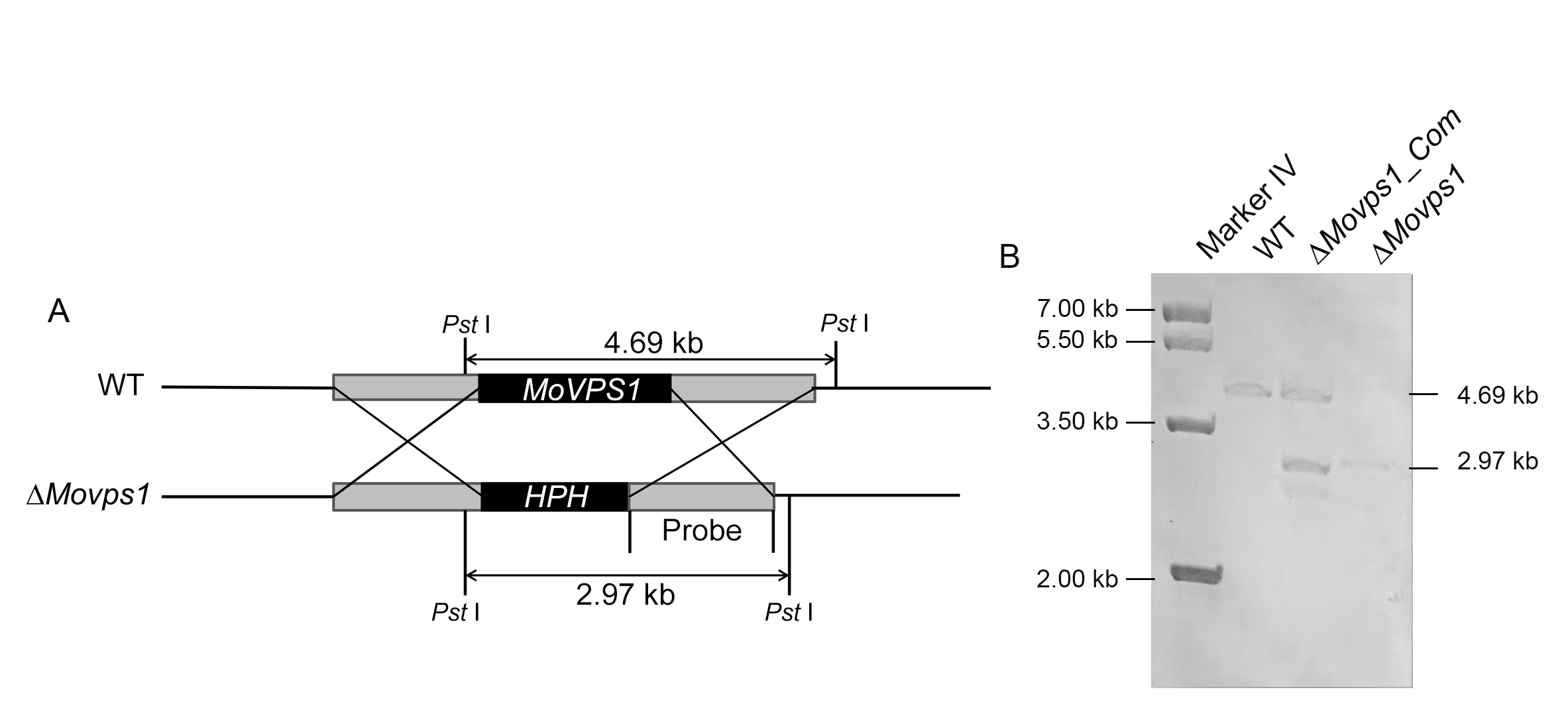

**Fig S6. Targeted gene deletion of *MoVPS1* in *M. oryzae***

Schematic representation of targeted gene disruption strategy used to generate Δ*Movps1* strains by homologous recombination. B. Southern blot result showing successful replacement of *MoVPS1* gene with hygromycin resistance gene in the Δ*Movps1* strains by single insertion, and successful reintroduction of the *MoVPS1* gene into the Δ*Movps1* strains.
